# Midbrain somatostatin-expressing cells control pain-suppression during defensive states

**DOI:** 10.1101/2022.01.20.476899

**Authors:** Nanci Winke, Frank Aby, Daniel Jercog, Coline Riffault, Rabia Bouali-Benazzouz, Juliette Viellard, Delphine Girard, Zoé Grivet, Marc Landry, Laia Castell, Emmanuel Valjent, Stephane Valerio, Pascal Fossat, Cyril Herry

**Affiliations:** Univ. Bordeaux, Neurocentre Magendie, U1215, 146 Rue Léo-Saignat, 33077 Bordeaux, France; INSERM, Neurocentre Magendie, U1215, 146 Rue Léo-Saignat, 33077 Bordeaux, France; Univ. Bordeaux, CNRS, Institute for Neurodegenerative Diseases, IMN, UMR 5293, F-33000 Bordeaux, France; Université de Montpellier, Institut de génomique fonctionnelle (IGF), CNRS, Inserm, 141 rue de la Cardonille, 34094 Montpellier, France; INM, Univ. Montpellier, Inserm, F-34094 Montpellier, France

## Abstract

In threatening situations, animals exhibit a broad range of behavioral and autonomic responses. As such, a crucial adaptive response is the inhibition of pain, which facilitates relevant defensive behaviors that promote survival. Whereas the structures and mechanisms involved in fear and pain behaviors are well documented, little is known about the precise neuronal mechanisms mediating the emotional regulation of endogenous pain-suppression. Here, we used a combination of behavioral, anatomical, optogenetic, and electrophysiological approaches to investigate, in male mice, the role of somatostatin-expressing cells in the ventrolateral periaqueductal gray matter (SST^+^ vlPAG cells) in the control of analgesia induced during defensive states. Our data indicate that optogenetic inhibition of SST^+^ vlPAG cells promotes analgesia irrespective of animal defensive state. In contrast, optogenetic activation of long-range SST^+^ vlPAG cells that project to the rostral ventromedial medulla (RVM) abolishes the analgesia mediated by fear behavior. Together, these results identify a novel circuit mechanism composed of long-range SST^+^ vlPAG cells projecting to the RVM that regulate analgesia elicited during defensive states.

## Introduction

During threatening situations, the inhibition of pain is an essential adaptive response that can be crucial to promote self-preservation^1–4^. The inhibition of pain responses to a stimulus that would typically elicit pain is defined as analgesia^5^. In addition, previous studies have described that a fearful event can reduce pain sensitivity, a phenomenon referred to as fear-conditioned analgesia (FCA)^6^. In particular, pharmacological inactivation of the basolateral amygdala abolishes this fear-modulated analgesia and is associated with increased immediate-early gene c-Fos expression in the central amygdala and periaqueductal gray matter (PAG)^7^. The PAG is a midbrain structure that receives many cortical and subcortical inputs from brain structures involved in fear processing^8–11^, projecting to the spinal cord primarily through a brainstem relay in the rostral ventromedial medulla (RVM)^12^, a key structure involved in nociceptive transmission^13,14^. Indeed, the ventrolateral part of the PAG (vlPAG) is crucial for the descending control of pain^13,15^ in the dorsal horn of the spinal cord (DH)^12^, and several reports indicate that electrical stimulation of the vlPAG selectively inhibits responses to noxious stimuli in a variety of pain test conditions^12^. Moreover, analgesia induced upon vlPAG stimulation is opioid-dependent^16–18^ and involves direct descending projections to the RVM and the DH^19–22^. Furthermore, previous research indicates that activation of the vlPAG induces analgesia through the recruitment of local GABAergic cells^23^, which is consistent with our understanding of the role of the GABAergic system in the descending control of pain^24,25^. Thus, the vlPAG is ideally positioned to mediate the emotional regulation of pain behavior, yet the underlying neuronal circuits and mechanisms are largely unknown. Here, we investigated the precise neural circuits within the vlPAG that mediate the suppression of pain behavior during threatening situations. We developed a unique behavioral paradigm where a cued-elicited defensive state promotes changes in pain sensitivity. Combining behavioral, anatomical, optogenetic, and electrophysiological approaches, we identified a novel midbrain to brainstem circuit mechanism composed of somatostatin positive (SST^+^) vlPAG cells projecting to the RVM that is critically involved in the suppression of pain behavior during conditioned fear states.

## Results

### Cued-fear conditioning elicits analgesia

To evaluate the contribution of specific vlPAG cell populations to the fear modulation of pain behavior, we developed a novel cued fear-conditioned analgesia (CFCA) paradigm, consisting of a pain sensitivity assay during the presentation of threat-predicting cues (**Figure 1a**). In this paradigm, an initially neutral stimulus (the conditioned stimulus, CS^+^) becomes associated with a footshock (the unconditioned stimulus, US), whereas another neutral stimulus serves as a control (CS^-^). After conditioning, re-exposure to the CS^+^ elicited freezing behavior, a behavioral readout of associative learned fear^26^ (**Figure 1b and Figure 1—figure supplement 1a**). Following the fear retrieval session, male mice were submitted to the pain sensitivity assay. Mice were placed on a Hot-Plate (HP) apparatus in which the floor plate temperature progressively increases while either CS^-^ or CS^+^ were concomitantly presented. Plate temperature increased, and CSs were continuously played until mice displayed a nociceptive (NC) response reflecting pain behavior (**Figure 1c**), defined by either jumping or hindpaw licking (**Figure 1d**). The detection of a NC response terminated the test session (**see Methods**).

**Figure 1.**
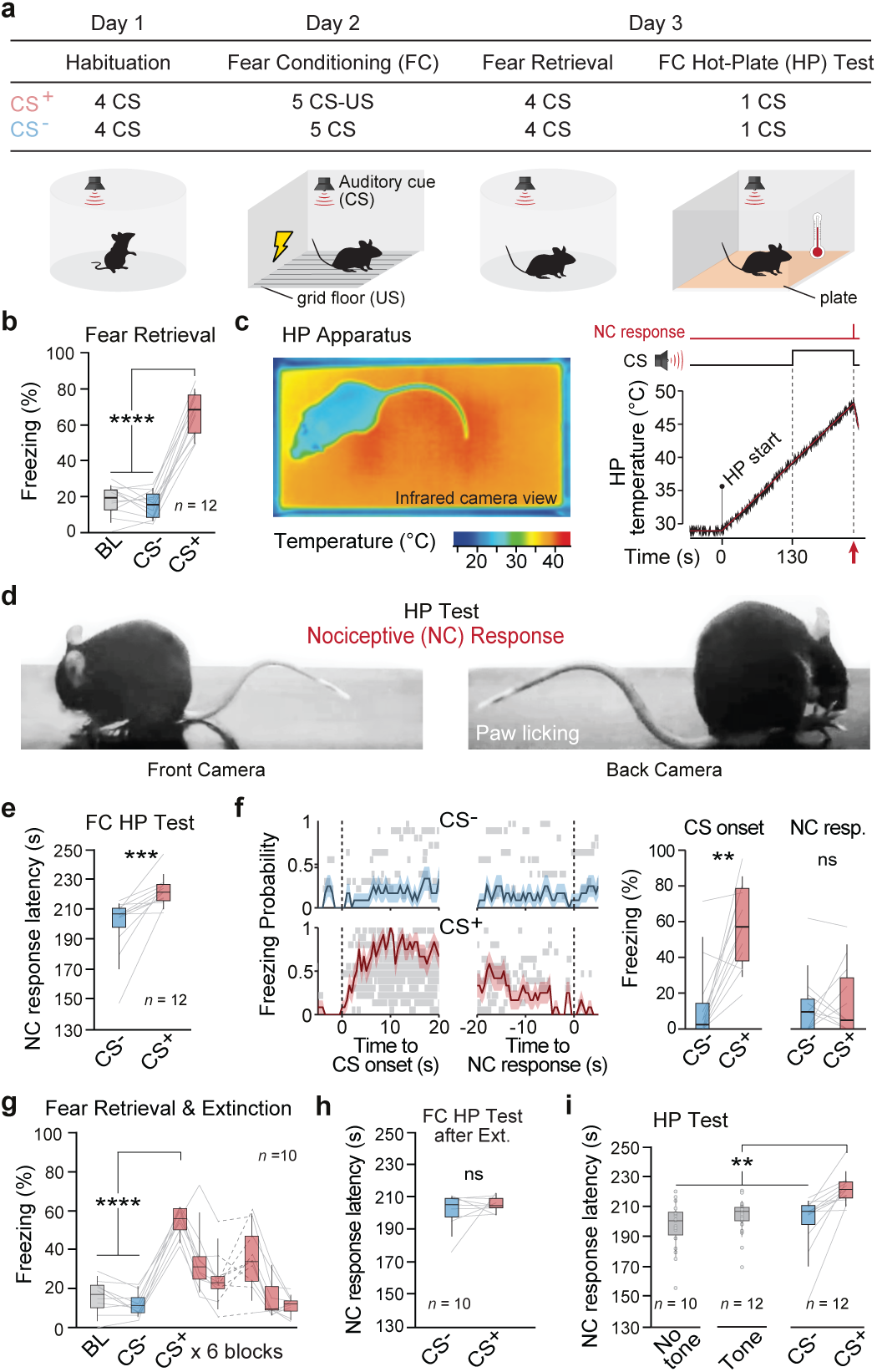
Fear modulation of pain behavior. **a.** Schematic of the setup and the CFCA paradigm. The paradigm consists of a discriminative auditory fear conditioning test followed by a pain sensitivity assay (for details, see Methods). **b.** Average freezing values for CS^+^ were higher than CS^-^ or baseline (BL) periods during retrieval (****, P < 0.0001, one-way RM ANOVA, n = 12 mice). **c.** Example of the infrared digital thermographic camera view of a mouse during the FC HP test. After 1 min of context acclimatization, the plate temperature steadily increases at 6 °C per min (HP start). CSs started 130 s after the HP start and continued until mice displayed an NC response (licking the hindpaw or jumping). **d.** Example of assessed hindpaw licking NC response. **e.** Latency of NC response on the FC HP test increased during CS^+^ compared to CS^-^(***, P = 0.0005, Wilcoxon matched-pairs signed rank test, n = 12 mice). **f. Left**: dynamics of freezing probability at CS onset and NC response for both CS^-^ and CS^+ (^freezing periods for each individual mouse are displayed by rows in grey; error bars display S.E.M; bin size = 0.5 s). **Right**: average freezing values during 10 s after CS onset and before NC response for CS^+^ were higher than CS^-^only at CS onset (***, P _onset_ = 0.0015, Wilcoxon matched-pairs signed rank test, n = 12 mice). **g.** Mean freezing values during fear extinction, 24 CS^+^ were presented across two separate extinction sessions (joined by dashed lines). Mice acquired the CS^+^-US association (1st CS^+^ block vs. BL/CS^-^****, P < 0.0001, Friedman test, n = 10 mice), followed by a rapid extinction (5th & 6th block of CS^+^ vs BL/CS-ns, P > 0.05, one-way RM ANOVA). **h.** After extinction, there was no difference in the NC response latency for both CSs (ns, P > 0.05, paired t-test n = 10 mice). **i.** Average NC response latencies for the different HP tests. *No tone* - mice submitted to the HP test without conditioning nor tone presentation. *Tone* - mice submitted to the HP test paired with unconditioned tone presentation. *CS^-^/CS^+^* - mice were submitted to the CFCA protocol. (No tone/Tone/CS^-^vs. CS^+^ **, P < 0.0001, Kruskal-Wallis test). For exact P values and test statistics on this and all subsequent figures, see Supplementary File 1.

We observed a significant delay in the NC response performance for CS^+^ compared to CS^-^presentations, reflecting that pain-suppression was selectively promoted by the threat-predicting CS (**Figure 1e and Figure 1—figure supplement 1b**). This increased NC latency during CS^+^ presentation could alternatively be explained by a competition between freezing and pain-related responses. However, freezing probability at the time of the NC response for CS^+^ was remarkably low and comparable to CS^-^levels, suggesting that CS^+^ increased NC response latency was not due to competing freezing responses (**Figure 1f, see Methods**). Instead, these results suggest that fear induced by CS^+^ presentations drives the increase in NC latency. Importantly, such states induced by our CFCA paradigm reflected fear associative processes (**Figure 1—figure supplement 2**). To demonstrate a causal link between cued-fear and analgesia, mice were submitted to an extinction procedure to devalue the CS^+^-US contingency and abolish the negative emotional state during CS^+^ presentations (**Figure 1g**). Under these conditions, CS^+^ presentation after extinction had no effects on NC response latency (**Figure 1h**). Furthermore, the absence of auditory cues, or the presence of auditory cues not associated with a threat, led to NC response latencies comparable to those observed during CS^-^presentations (**Figure 1i**). Finally, previous studies indicate that fear can lead to vasoconstriction, which could result in a redirection of the blood flow to the skeletal musculature, decreasing the temperature in the extremities of the body^27^. However, monitoring the back and tail temperatures of mice submitted to the HP test did not reveal any differences between the CSs presentations, suggesting that the increased latency was not due to fear-induced vasoconstriction (**Figure 1—figure supplement 1c-e)**. Overall, our results indicate that the presence of a conditioned threat-predicting cue can selectively promote analgesia.

### Photostimulation of SST^+^ vlPAG cells alters pain and fear behavior

To identify the cell types in the vlPAG mediating the analgesic effect in the CFCA task, we focused on GABAergic interneurons, whose activity is suggested to be inhibited during analgesia through µ-opioid receptor-dependent mechanisms^12,25^. Using single-molecule fluorescent in situ hybridization (smFISH, **see Methods**), we confirmed that the majority of vlPAG *Sst* mRNAs were expressed in inhibitory neurons (**Figure 1—figure supplement 3**). As such, since somatostatin-expressing interneurons (SST) are the most abundant interneuron class in the vlPAG^28^, we focused on this cell population. Thus, to evaluate the contribution of SST^+^ vlPAG cells in CFCA, *Sst-IRES-Cre* mice were injected in the vlPAG with a Cre-dependent AAV expressing the Channelrhodopsin (ChR2), Archaerhodopsin (ArchT), or GFP while optic fibers were bilaterally implanted targeting this area (**Figure 2a**, **Figure 2—figure supplement 1**). Next, mice were submitted to the CFCA task while photostimulating SST^+^ cells in the vlPAG (**Figure 2b**; **see Methods**). After fear retrieval (**Figure 2c, e**; **Figure 2—figure supplement 2a-d**), the optogenetic activation of SST^+^ vlPAG cells during the HP test had no effect on CS^-^presentation, whereas the same manipulation performed during CS^+^ blocked the increase in NC response latency compared to GFP controls (**Figure 2d and Figure 2—figure supplement 3a**). In contrast, optogenetic inhibition of SST^+^ vlPAG cells increased NC response latency for both CS^-^ and CS^+^ conditions compared to GFP controls (**Figure 2f and Figure 2—figure supplement 3b**). Importantly, these changes in NC response latencies were not due to motor or aversive effects of the optogenetic manipulations, since photoactivation of SST^+^ vlPAG cells did not induce changes in distance traveled in an open field nor place aversion in a real-time place preference test (**Figure 2—figure supplement 3c-f**). In addition, we did not observe changes in fear learning or expression when the footshock was replaced or paired with photoactivation of the SST^+^ vlPAG cells during the fear conditioning procedure (**Figure 2—figure supplement 4**).

**Figure 2.**
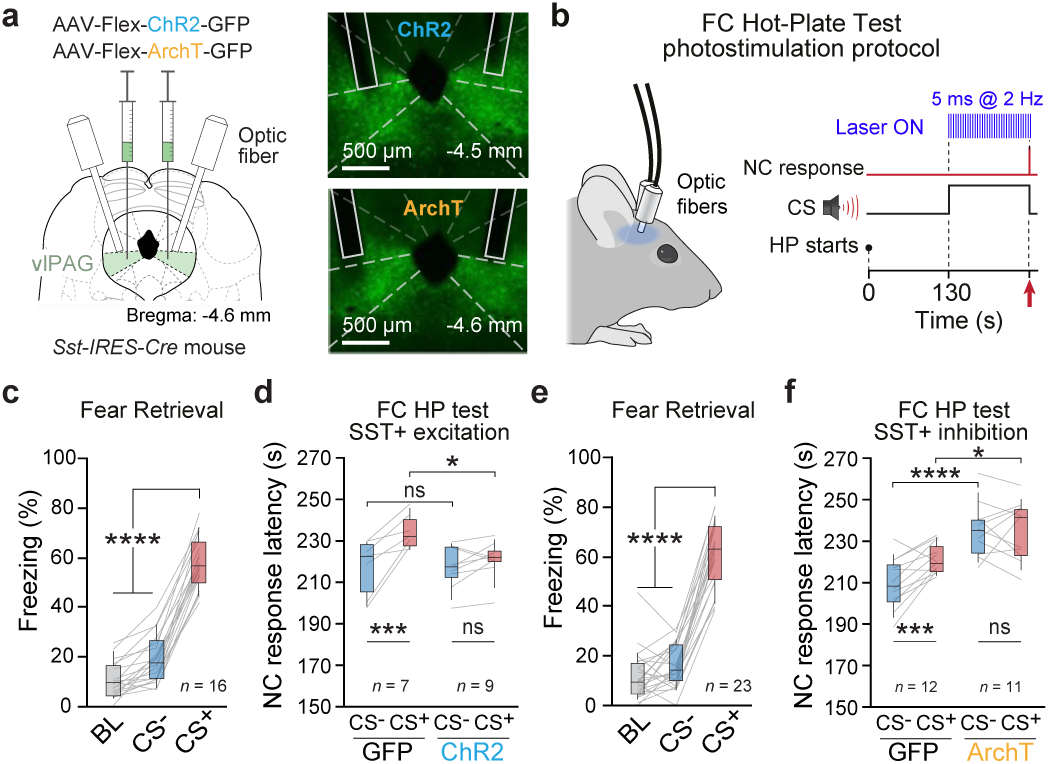
SST^+^ cells in vlPAG mediate the fear modulation of pain behavior. **a**. *Sst-IRES-Cre* mice were bilaterally injected with viruses that expressed ChR2 (right, upper), ArchT (right, bottom), or GFP (control) in SST^+^ vlPAG cells. **b**. Schematics of the stimulation protocol used for photoactivation of the SST^+^ vlPAG cells: blue light delivered at 2 Hz with 5 ms pulse duration during CSs presentation until NC response. For photoinhibition experiments, green light was delivered continuously. **c, e**. Average freezing values during retrieval for ChR2-(**c**) and ArchT-infected mice (**e**) and their respective GFP controls. The opsin and respective control groups were pooled together because no difference was found in the conditioning level (see Figure 2—figure supplement 2). Average freezing values during CS^+^ were higher than CS^-^ or baseline (BL) periods (****, P < 0.001, one-way RM ANOVA, (**c**) n = 16 mice and (**e**) n = 23 mice). **d**. Photoactivation of SST^+^ vlPAG cells abolished the analgesic effect induced by fear (*, P = 0.0112, opsin x CS - two-way RM ANOVA, n = 7 GFP, n = 9 ChR2). The NC response latency for the CS^+^ was significantly different between the ChR2 and GFP groups (*, P = 0.0344, Bonferroni post-hoc test). For the ChR2 group, the NC response latency during CS^+^ was equivalent to the CS^-^(ns, P = 0.3876, Bonferroni post-hoc test). On the contrary, NC response latency between the CSs differed in the GFP group (***, P = 0.0003, Bonferroni post-hoc test). **f**. Photoinhibition of SST^+^ vlPAG cells increased the analgesic effect for the ArchT group when compared to the GFP (*, P = 0.0037, opsin x CS - two-way RM ANOVA, n = 12 GFP, n = 11 ArchT). The NC response latency for the CS^-^ and CS^+^ was significantly different between the ArchT and GFP group (CS^-^: ****, P < 0.0001, Bonferroni post-hoc test; CS^+^: *, P = 0.0265, Bonferroni post-hoc test). For the GFP group, the latency of NC response was higher for the CS^+^ trials when compared to the CS^-^trials (***, P = 0.0003, Bonferroni post-hoc test), yet this was not the case for the ArchT group (ns, P > 0.999, Bonferroni post-hoc test).

Altogether, these experiments confirmed that SST^+^ vlPAG activation did not promote pain responses and instead suggested that the SST^+^ vlPAG cells are key elements of an endogenous pain-suppression circuit. Additionally, the optogenetic effect observed during CFCA was not due to alteration of somatostatin levels in our homozygous *Sst-IRES-Cre* mice^29^, as we observed the same effect in heterozygous *Sst-IRES-Cre* mice (**Figure 2—figure supplement 5a-d**). Finally, in contrast with STT^+^ vlPAG cells, the optogenetic inhibition of another important class of vlPAG inhibitory cells expressing the vasoactive intestinal peptide (VIP^+^) did not change the NC response latencies during CS^-^ or CS^+^ presentations (**Figure 2—figure supplement 5e-h, Figure 2—figure supplement 6**).

Because our CFCA paradigm is dependent on fear associative processes (**Figure 1 and Figure 1—figure supplements 1 and 2**), it is possible that the optogenetic manipulation of SST^+^ vlPAG cells may have altered the expression of fear behavior. To control for this possibility, we optogenetically activated or inhibited SST^+^ vlPAG cells during a fear retrieval session following auditory fear conditioning (**Figure 3a, b**). Our data indicate that whereas optogenetic inhibition of SST^+^ vlPAG cells had no effect on freezing behavior relative to GFP controls, their optogenetic activation reduced freezing levels (**Figure 3c, d**). Altogether, our data show that the local SST^+^ vlPAG network can modulate both fear expression and the analgesia promoted during defensive states.

**Figure 3.**
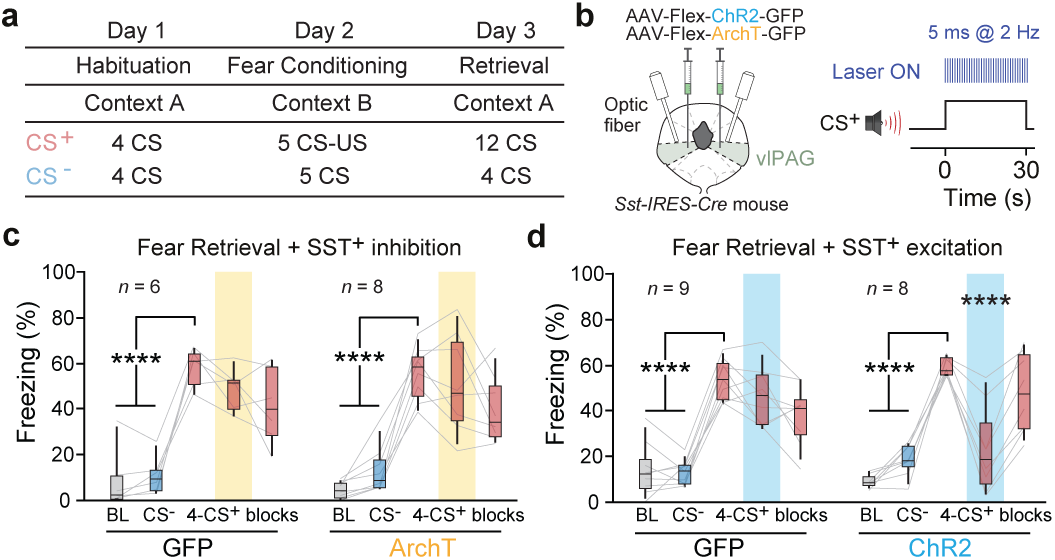
Activation of SST^+^ vlPAG cells reduced fear expression. **a**. Protocol for optogenetic manipulation during fear retrieval. Days 1 and 2 were done as described previously for the CFCA paradigm. During retrieval, there were 12 CS^+^ presentations divided into three blocks. **b**. The stimulation protocol used to photoactivate the SST^+^ vlPAG cells. The optogenetic manipulation was performed during the 2nd block of the CS^+^ presentation (analogously to CFCA manipulations, **Figure 2b**). **c**. Light inhibition of SST^+^ vlPAG cells did not modulate freezing levels (ArchT ns, P = 0.9396, opsin x CS two-way RM ANOVA, n = 6, GFP, n = 8). **d**. Photoactivation of the SST^+^ vlPAG cells had no effect on the GFP group, but it transiently decreased the freezing levels for the ChR2 group (***, P < 0.0001, opsin x CS two-way RM ANOVA, n = 9 GFP, n = 8 ChR2).

### Photostimulation of SST^+^ vlPAG cells regulates spinal cord nociceptive transmission

Consistent with our observations, previous studies found that the vlPAG is involved in fear-conditioned analgesia^7^. It is conceivable that the SST^+^ vlPAG cells are indeed part of a circuit involved in the fear regulation of endogenous pain-suppression. To evaluate the effect of the manipulation of SST^+^ vlPAG cells on analgesia independently of the expression of fear behavior, we evaluated the impact of SST^+^ vlPAG cells manipulation on the control of nociceptive transmission directly at the level of the spinal cord DH. Therefore, we performed extracellular local field potential and single-unit recordings in the spinal cord DH during optogenetic stimulation of SST^+^ vlPAG cells in anesthetized mice (**Figure 4a**). Nociceptive stimulation induces early and late neuronal responses at the level of the spinal cord DH^30^ (A-comprising non-nociceptive Aβ and nociceptive Aδ and C-fiber mediated, respectively; **Figure 4a**). While optogenetic activation of SST^+^ vlPAG cells during suprathreshold electrical stimulation of the paw increased the magnitude of late nociceptive field potentials, the optogenetic inhibition reduced it (**Figure 4b, c**). To determine whether this effect was specific to the nociceptive network, we recorded wide dynamic range (WDR) cells in the DH, which receive both tactile (Aβ) and nociceptive (Aδ and C) information^30^. We observed that photoactivation of SST^+^ vlPAG cells increases both fast nociceptive (Aδ-fiber) and slow nociceptive (C-fiber) responses without affecting non-nociceptive (Aβ-fiber) responses (**Figure 4d**). Moreover, optogenetic inhibition of SST^+^ vlPAG cells specifically inhibited nociceptive responses, whereas non-nociceptive responses (Aβ-fiber) remained unaffected (**Figure 4e**). In addition, we performed subliminal electrical stimulations that did not elicit nociceptive neuronal responses. In this condition, we observed that removing analgesia by activating SST^+^ vlPAG cells elicited WDR responses at latencies corresponding to nociceptive Aδ- and C-fibers (**Figure 4f**). Finally, the optogenetic activation of SST^+^ vlPAG cells had no effect on purely tactile fast latency spinal neurons (**Figure 4—figure supplement 1**). Consistent with our optogenetic manipulations during the CFCA task (**Figure 2**), these data suggest that, under analgesia promoted by isoflurane, activation and inhibition of SST^+^ vlPAG cells increased and decreased nociceptive transmission, respectively, by specifically promoting pronociceptive and antinociceptive responses in the spinal cord DH.

**Figure 4.**
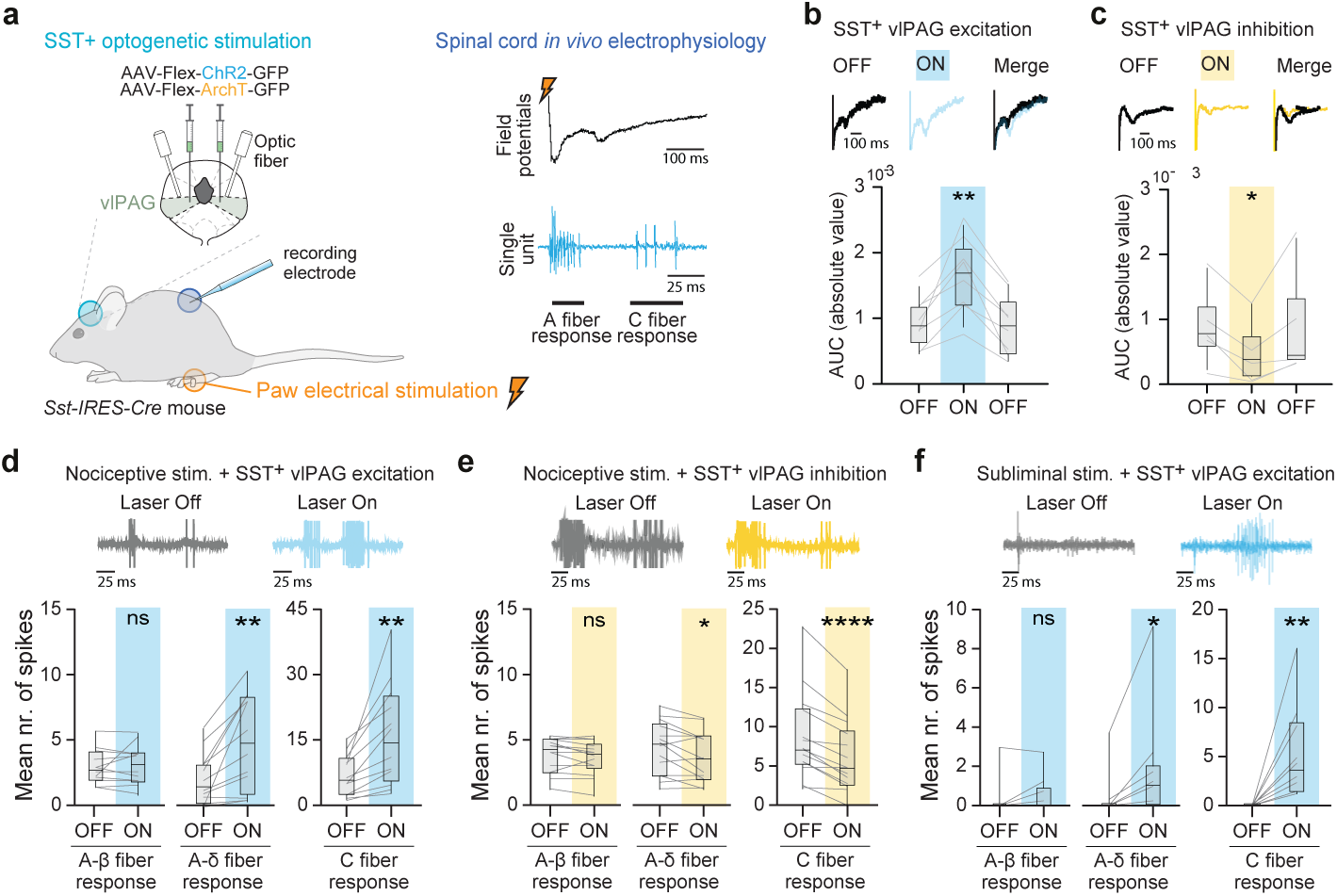
Manipulation of SST^+^ vlPAG cells alters spinal cord-related pain signals. **a.** Schematics of *in vivo* anesthetized experiments. Extracellular local field potential and single-unit recordings in the spinal cord DH were performed during optogenetic stimulation of SST^+^ vlPAG cells with concomitant noxious electrical paw stimulation**. b.** Example of average nociceptive field potentials in the lumbar SC before (OFF), during (ON), and after (OFF) photoactivation of SST^+^ vlPAG cells (top). Photoactivation induces a significant increase in the nociceptive fields (bottom;**, P = 0.0001, one-way RM ANOVA, n = 8 mice). **c.** Analogous to panel b, but for photoinhibition of SST^+^ vlPAG cells. Photoinhibition induces a significant decrease in the nociceptive fields (bottom; *, P = 0.0275, one-way RM ANOVA, n = 5 mice). **d.** Example of WDR single-unit activity before and during photoactivation of SST^+^ vlPAG cells (**top**). Photoactivation of SST^+^ vlPAG cells induces a significant and global increase in WDR response to both Aδ- and C-mediated nociceptive fibers (**bottom**; **, P_Aδ fiber = 0.0022, paired t-test, n = 12 cells;_ **, P_C fiber = 0.0033, paired t-test, n = 12 cells)_. **e.** Analogous to panel d, but for photoinhibition of SST^+^ vlPAG cells. Photoinhibition induces a significant and specific inhibition of WDR response to Aδ- and C-fibers (**bottom**; *, P_Aδ fiber =_ 0.0241, ****, P_C fiber <_ 0.0001, n = 14 cells, paired t-test). **f.** Example traces of single-unit recordings of WDR cells with subthreshold electrical stimulation accompanied by photoactivation of SST^+^ vlPAG cells (**top**). The photoactivation of SST^+^ vlPAG cells elicits WDR response to A- and C-mediated peripheral fibers (**bottom**; *, P_Aδ fiber =_ 0.0156, n = 10 cells; **, P_C fiber =_ 0.0020, n = 10 cells, Wilcoxon matched-pairs signed rank test).

### SST^+^ vlPAG-RVM-spinal cord pathway activation abolishes analgesia induced during defensive states

We next evaluated whether SST^+^ vlPAG cells mediate their analgesia modulation by contacting WDR cells directly in the spinal cord or through the rostro ventromedial medulla (RVM), a structure known to receive direct projections from vlPAG and mediate pain modulation^19,20^ (**Figure 5a**). Anatomical tracing revealed large labeling of SST^+^ fibers in RVM (**Figure 5b**) and only sparse labeling within the spinal cord (SC; data not shown). By combining fluorogold retrolabeling from the SC and GFP anterograde labeling from SST^+^ vlPAG cells (**Figure 5c**), we observed RVM cells projecting to the DH with close apposition of SST^+^ putative boutons (**Figure 5d**), suggesting that SST^+^ vlPAG cells project to the RVM and contact DH-projecting RVM cells. Optogenetic activation of SST descending fibers above the SC has no effect on WDR neural activity upon paw electrical stimulation, suggesting that SST^+^ vlPAG long-range projections to SC did not play a major role in endogenous pain-suppression (**Figure 5e**). In contrast, photoactivation of SST^+^ vlPAG inputs to the RVM increased both nociceptive (Aδ and C-fiber) responses (**Figure 5f**, **Figure 5—figure supplement 1**). Altogether, these data revealed that the SST^+^ vlPAG-RVM-DH pathway mediates endogenous pain-suppression.

**Figure 5.**
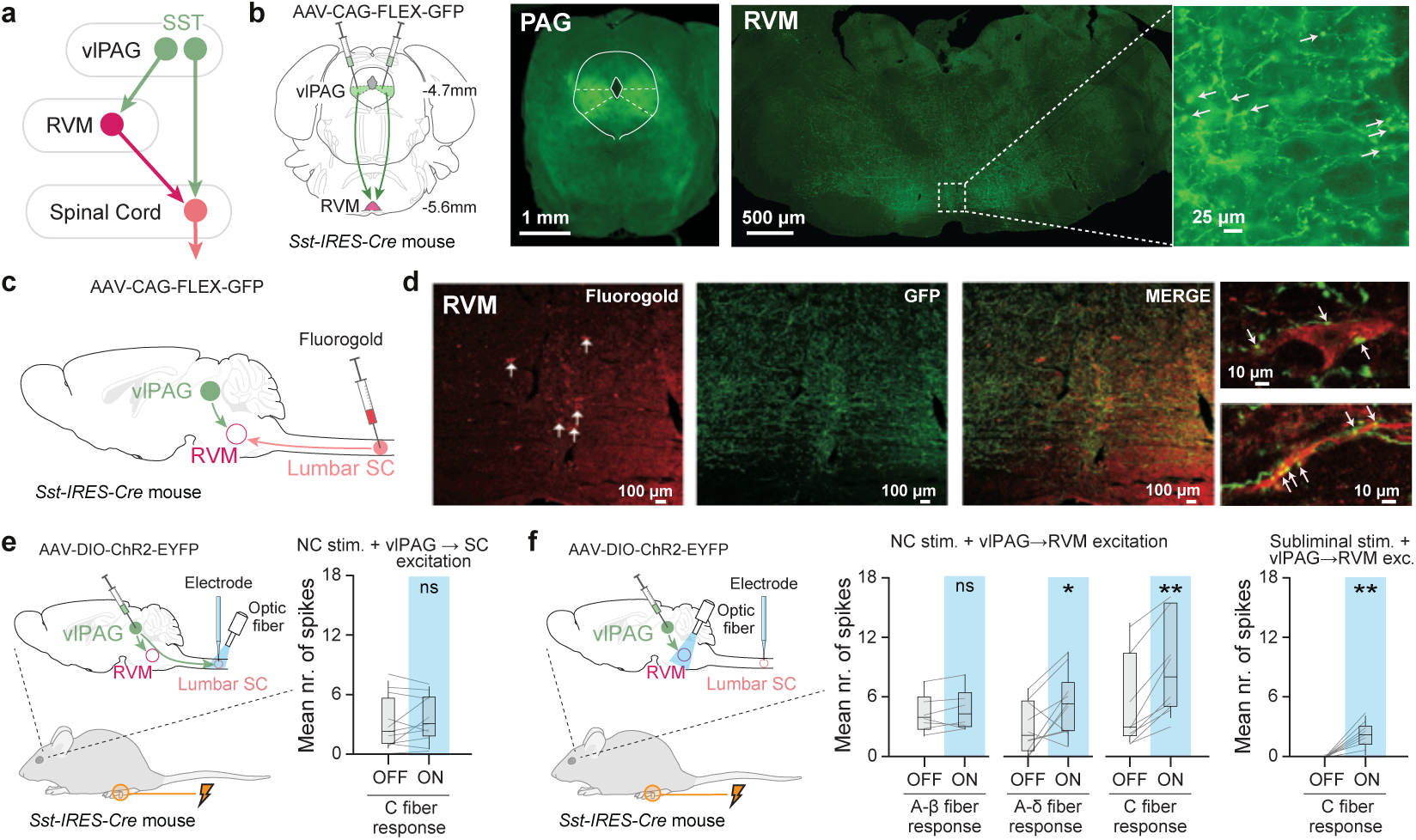
SST^+^ vlPAG-RVM-DH pathway activation removes analgesia. **a**. Schematics of possible SST^+^ vlPAG circuits mediating analgesia by long-range projections directly to the SC or, alternatively, by projecting to the SC via the rostral ventromedial medulla (RVM). **b**. *Sst-IRES-Cre* mice were injected with an anterograde AAV Cre-dependent GFP virus in the vlPAG (left). Example of SST fibers labeling at the level of the RVM (middle). Higher magnification (right) reveals putative axonic buttons in the RVM (examples indicated by white arrows). **c**. *Sst-IRES-Cre* mice were injected concomitantly with anterograde AAV Cre-dependent GFP virus in the vlPAG and retrograde fluorogold in the lumbar DH of the SC. **d**. Fluorogold positive cells (red) cross SST^+^ vlPAG fibers in the RVM (green). Higher magnification (**right**) shows close contacts between the putative SST^+^ button and fluorogold^+^ cells or fibers (white arrows). **e**. *Sst-IRES-Cre* mice were injected in the vlPAG with AAV Cre-dependent ChR2 virus and an optic fiber placed above the lumbar SC. Single-unit recordings of WDR cells while photoactivation of the SST^+^ vlPAG SC fibers and electrically stimulating the paw in anesthetized mice (**left**). Photoactivation of vlPAG projections to SC did not affect nociceptive transmission (**right**; ns, P = 0.3683, paired t-test, n = 11 cells). **f**. Same experimental design as in panel e, except that the optic fiber was placed above the RVM (left). Photoactivation of SST^+^ vlPAG inputs to the RVM induces a significant increase in WDR response to both Aδ- and C-mediated nociceptive fiber stimulation (**middle**; *, P_A*δ* fiber =_ 0.0346, Paired t-test, n = 10 cells; **, P_C fiber =_ 0.0020, Wilcoxon matched-pairs signed rank test, n = 10 cells). Removal of analgesia by activation of SST^+^ vlPAG-RVM induces a significant increase in WDR response to C-fiber responses (right; **, P = 0.0078, Wilcoxon matched-pairs signed rank test, n = 8 cells).

We next addressed whether the SST^+^ vlPAG-RVM-DH pathway was specific to mediating endogenous pain-suppression induced by defensive states. Thus, *Sst-IRES-Cre* mice were injected in the vlPAG with a Cre-dependent AAV expressing the ChR2 or GFP, and a single optic fiber was implanted above the RVM (**Figure 6a**). While photoactivation of the SST^+^ vlPAG-RVM pathway promoted nociception (**Figure 5f**), it did not impact fear expression as freezing levels remained unchanged (**Figure 6b**). Consistently, activation of the SST^+^ vlPAG-RVM pathway during CFCA had no impact on CS^-^presentation, whereas the same manipulation performed during CS^+^ blocked the increase in NC response latency compared to GFP controls (**Figure 6c, d**). Overall, these data suggest the existence of two populations of SST^+^ vlPAG cells mediating fear expression and analgesia induced by defensive states. (**Figure 6e**). Altogether, our results indicate that the SST^+^ vlPAG-RVM-DH pathway is critically involved in the analgesia induced during defensive states.

**Figure 6.**
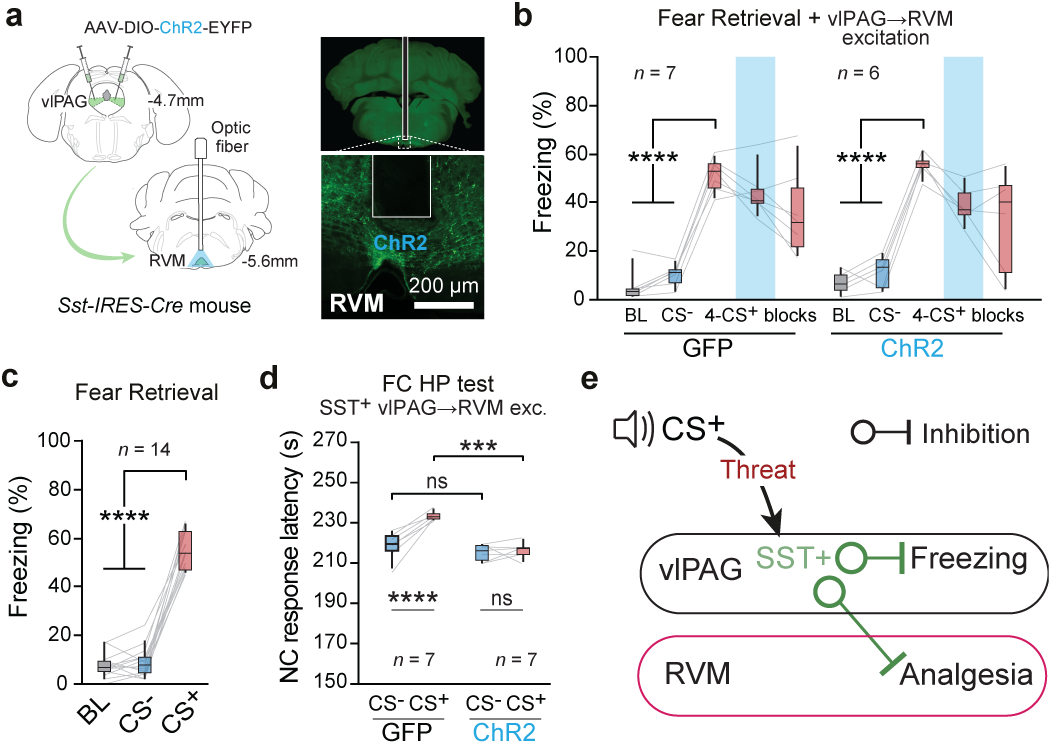
SST^+^ vlPAG-RVM-SC pathway activation abolishes analgesia induced by CFCA. **a.** *Sst-IRES-Cre* mice were bilaterally injected with an AAV expressing ChR2 or GFP in the vlPAG, and optic fibers were implanted above the RVM (left). Representative example of SST^+^ vlPAG terminals in the RVM (right). **b.** Photoactivation of SST^+^ vlPAG-RVM pathway did not modulate freezing levels (ns, P = 0.6443, opsin x CS two-way RM ANOVA; CS^+^ 2nd block, GFP vs. ChR2 ns, P > 0.9999, Bonferroni post-hoc test, n = 7 GFP, n = 6 ChR2). **c.** Average freezing values during retrieval for pooled ChR2 and GFP-infected mice (groups were pooled together since no difference was found in the conditioning level). Average freezing values during CS^+^ were higher than CS^-^ or baseline (BL) periods (****, P < 0.0001, one-way RM ANOVA, n = 14 mice). **d.** Photoactivation of SST^+^ vlPAG-RVM pathway abolished the analgesic effect modulated by fear (***, P = 0.0008, opsin x CS - two-way RM ANOVA, n = 7 GFP, n = 7 ChR2). For the ChR2 group, the NC response latency during CS^+^ was equivalent to the CS^-^(ns, P > 0.9999, Bonferroni post-hoc test). On the contrary, the NC response latency between the CSs differed in the GFP group (****, P < 0.0001, Bonferroni post-hoc test). **e**. Working model schematics for SST^+^ control analgesia induced during defensive states. Activation of SST^+^ cells in the vlPAG is associated with the inhibition of fear-induced freezing behavior. Selective activation of RVM projecting SST^+^ vlPAG cells is associated with inhibition of analgesia induced during defensive states.

## Discussion

In this study, we established that SST^+^ vlPAG cells control analgesia during defensive states in male mice. Endogenous pain-suppression was mediated by long-range SST^+^ vlPAG cells projecting to the RVM, which, in turn, transmitted the pain signal to the DH of the spinal cord (**Figures 2 and 5**). Finally, our data indicated that SST^+^ vlPAG cells can regulate both fear expression and the analgesia promoted during defensive states, and that the expression of these two behaviors can be segregated by precisely targeting the SST^+^ vlPAG-RVM pathway (**Figure 6**). Together, these results identify a novel midbrain to brainstem circuit mechanism composed of long-range SST^+^ vlPAG cells projecting to the RVM that regulate analgesia elicited during defensive states. Pain mechanisms and their modulation are known to exhibit sex-dependent differences^31^, and thus we cannot exclude the possibility that the circuits described here may differ in females. Nevertheless, recent studies of RVM-spinal cord descending control suggest broadly similar organization and function across sexes^32,33^. Direct experimental investigation in female animals will be required to determine the extent to which the present findings generalize across sexes.

Our observations of the involvement of vlPAG in our cued-based fear conditioned analgesia paradigm are consistent with previous studies of contextual-based FCA involving this region^7^. In such FCA studies^6^, animals undergo contextual fear conditioning followed by pain behavior testing in the same conditioning context, thereby limiting the temporally precise elicitation of aversive states. Instead, our behavioral paradigm goes beyond these limitations by promoting defensive states through a cued conditioned stimulus, advancing toward temporally specific manipulations such as optogenetics, and allowing us to study the phenomena with a within-subject experimental design through subsequent CS^+/-^presentations. Nevertheless, since the circuits involved in contextual and cued fear may differ^30,34^, this might also imply different circuits involved in the emotional modulation of pain-suppression behavior. To our knowledge, this is the first demonstration of a cued fear-conditioned analgesia model in rodents. Addressing potential differences in the circuitry involved in contextual and cued fear-induced analgesia will require further investigation.

Our observation that photoactivation of SST^+^ vlPAG cells not only abolished analgesia during defensive states but also decreased freezing expression has far-reaching functional consequences. Two independent previous studies found that long-range inhibitory inputs from the central medial amygdala contact inhibitory cells within the vlPAG, implicated in different roles: the modulation of fear behavior^23^ and nociceptive transmission^35^. Notably, both studies describe the involvement of vlPAG inhibitory cells in a disinhibitory circuit mechanism of long-range projecting excitatory cells as the underlying neural substrates responsible for such modulations^23,35^. We observed that SST^+^ cells are the most abundant inhibitory cell type in the vlPAG (**Figure 1—figure supplement 3**). Thus, it is conceivable that the SST^+^ vlPAG cells are indeed part of a circuit involved in the emotional regulation of endogenous pain-suppression. Notably, global manipulation of SST^+^ vlPAG cells cannot disentangle their role in analgesia and fear behavior at the local level (**Figures 2 and 3**). Because the studies mentioned above exclusively addressed the contribution of vlPAG circuits to either pain^35^ or fear^23^ behavior, these studies may have been oblivious to alternative behavioral readout interpretations. Here, we demonstrated that SST^+^ vlPAG cells projecting to the RVM modulate analgesia induced during defensive states without impacting the behavioral expression of fear (**Figure 6**). Thus, our results suggest that fear and analgesia signals rely on overlapping neuronal circuits and that the discrimination between these two states could be mediated within the vlPAG network. Altogether, our data are consistent with the existence of different populations of SST^+^ cells within the vlPAG that mediate fear behavior and analgesic information, defined by their output connectivity.

The lateral inhibition model of analgesia within the PAG relies on the activation of PAG excitatory cells projecting to the RVM. More precisely, in this model, analgesia is thought to occur through an opioid-dependent inhibition of GABAergic cells, ultimately disinhibiting excitatory cells projecting to the RVM^12,20,21^. Our data indicate that, in addition to this disinhibitory mechanism, analgesia is also mediated by the inhibition of a direct long-range SST^+^ neurons projection onto RVM cells projecting to the DH (**Figure 6**). In line with this, inhibition of vlPAG SST cells induced analgesia in our experiments, consistent with previous findings obtained using chemogenetic approaches^36^. However, important differences emerge when comparing our results with Zhang et al.^34^. In that study, activation of vlPAG SST cells produced hyperalgesia, whereas in our study, activation of SST^+^ vlPAG cells did not induce hyperalgesia but instead suppressed fear-conditioned analgesia. One key distinction between the two studies is the experimental context: Zhang et al. examined this circuit in a neuropathic pain model, whereas our work focuses on acute nociceptive processing and fear-dependent modulation of pain. Thus, the functional contribution of SST^+^ vlPAG cells may depend on the behavioral and pathological state, with potentially distinct roles in chronic versus acute pain conditions^35^. In addition, methodological differences, such as our nociceptive test that uses a slow temperature ramp, may impose a floor effect that limits the detection of thermal hyperalgesia. Our results are therefore consistent with the parallel inhibition-excitation model, in which inhibitory and excitatory cells form two distinct, parallel descending pathways for pain modulation^24^. Indeed, previous studies have identified inhibitory projections within the PAG–RVM-spinal cord dorsal horn neuraxis^37^, and our results suggest that SST^+^ vlPAG cells contribute to analgesia within this broader framework. At the same time, our data indicate that vlPAG SST^+^ cells are not exclusively inhibitory, as approximately one third co-localize with excitatory markers, and recent work has shown that excitatory SST^+^ vlPAG cells project to the RVM^36^. Together, these observations raise the possibility that distinct SST^+^ subpopulations contribute differently to descending pain control, with long-range vlPAG SST^+^ cells participating in the excitatory pathway projecting throughout the PAG–RVM-spinal cord dorsal horn neuraxis, and local GABAergic vlPAG SST^+^ cells contributing to local circuit dynamics related to fear response, such as freezing^23^. We identified a novel midbrain to brainstem circuit mechanism composed of SST^+^ vlPAG cells that project to the RVM, thereby regulating analgesia during defensive states. Substantial progress has been made in outlining the midbrain circuitry of the endogenous pain-suppression pathway^12,24,26^. However, only more recently have high cortical areas been shown to be involved in analgesia^7,38^. Previous reports describe connections between CeL and vlPAG inhibitory neurons in the modulation of pain behavior^35,39^. Future work is required to elucidate whether the exact circuit that regulates analgesia is mediated by different negative emotional states, such as fear, depression, or stress.

Recent human reports reveal strong comorbidity between mood and anxiety disorders with and pain-related disorders^40^, suggesting that the deregulation of the circuits mediating the interplay between emotional and pain behavior could be at the core of such psychiatric conditions. Comprehensive knowledge about the circuits mediating the emotional regulation of pain behavior is urgently needed. We identified an SST^+^ cell type- and pathway-specific involvement in the mechanisms that mediate the emotion-dependent regulation of pain behavior. Thus, these findings may empower the development of more effective analgesic therapies by the improvement of further refined pharmacological targets.

## Methods

### Subject details

We used either male C57BL/6J mice (Janvier), heterozygous or homozygous *Sst-IRES-Cre* mice (Jackson laboratory, Stock # 013044), or heterozygous *Vip-IRES-Cre* mice (Jackson Laboratory, Stock # 010908) age 8-14 weeks, that were individually housed under a 12 h light-dark cycle and provided with food and water ad libitum. All procedures were performed in accordance with standard ethical guidelines (European Communities Directive 86/60-EEC) and were approved by the committee on Animal Health and Care of Institut National de la Santé et de la Recherche Médicale and the French Ministry of Agriculture and Forestry (agreement #A3312001).

### Behavioral apparatus

***Cued fear-conditioned analgesia*** *task* was performed in three different contexts (**Figure 1a**). Context A was used for *Habituation* and *Retrieval* and consisted of a plexiglass cylinder (25 x 24 cm diameter) with a grey, smooth plastic floor and house lights (16 lux). Context B was used for *Fear conditioning* and consisted of a square plexiglass (25 x 40 cm) with a grid floor connected to a shocker (Coulbourn Instruments) and brighter house lights (40 lux). A total of 5 scrambled footshocks of 1 s duration and intensity of 0.8 mA were delivered via the grid floor and served as the unconditioned stimulus (US). Contexts A and B were cleaned, respectively, with 70% ethanol or 1 % acetic acid between different mice. Both contexts contained infrared beam detection that automatically scored freezing periods, determined by no change in infrared beam crossing for at least 2 s.

Context C was used for the *FC Hot Plate (HP) test*. A steady increase in temperature was controlled by the Incremental Hot/Cold Plate Analgesia Meter (IITC) device. To restrain the mice’s movement, the HP apparatus had a testing surface (plate) enclosed in a square plexiglass surface (20.3 x 10 x 20.5 cm) and house lights (100 lux). The mice’s temperature and surroundings were recorded using an infrared digital thermographic camera (Testo 885) placed ∼ 50 cm above the testing surface. The thermal camera had a spatial resolution of 320 × 240 pixels, a sampling rate of 25 Hz, and thermal sensitivity of 0.03 °C at 30 °C. The testing surface of context C was cleaned with water between different mice. All three contexts were enclosed in an acoustic foam isolated box with speakers mounted on the top of each compartment. The auditory conditioned stimulus (CS) consists of either 7.5 kHz or white-noise 50 ms pips at 1 Hz, repeated 27 times, with 2 ms rise and fall, and a sound pressure level of 80 dB.

***Open field*** *task* was performed in a square plexiglass arena (36 x 36 x 25cm). A LED mounted on the top-right side of the arena signaled the start and end of different epochs for offline analyses. A video camera recorded from above the arena at 30 fps for offline video-tracking purposes.

***Real-Time Place Preference (RTPP)*** *task* was performed in a shuttlebox consisting of a plexiglass box (40 x 10 x 30 cm) with a floor grid, where a small plastic hurdle (1 cm height) divided the arena into two equal compartments, while infrared beam detection automatically monitored the mice shuttling between compartments (Imetronic). A video camera recorded from above the arena at 30 fps for offline video-tracking purposes.

For both the open field and RTPP tasks, a free user video-tracking software (idTracker: Tracking individuals in a group by automatic identification of unmarked animals) together with in-house codes in Matlab (The MathWorks, Inc., Natick, MA, USA) was used to analyze each condition.

### Behavioral paradigm

#### Cued fear-conditioned analgesia (CFCA)

C57BL/6J mice (n = 12) were habituated to the context and tones (day 1). Four white-noise (CS^-^) and four 7.5 kHz (CS^+^) tones were presented sequentially and without US reinforcement. In Fear conditioning (day 2), five CS^-^ and five CS^+^ were presented in an intermingled fashion. The CS^+^ presentations were paired with a mild footshock (US) at tone offset, whereas the CS^-^was never reinforced. The Fear retrieval session (Day 3) was done 24h after conditioning and in the same context as the habituation session. As in habituation, four CS^-^ and four CS^+^ were presented sequentially and without US reinforcement (**Figure 1a**).

Since the focus of this study was the emotional modulation of pain sensitivity, it was compulsory to evaluate the associative fear levels before measuring their impact on pain sensitivity. Two indices were computed: the discrimination index (DI), to assess the discrimination between CS^-^ and CS^+^, and the conditioning index (CI), which indicated the level of freezing to the tone predicting the US. These indexes were calculated as follows: 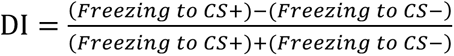 and CI = DI × (Freezing to CS^+^). Based on preliminary data, mice were only submitted to the FC HP test if DI ≥ 0.4 & CI ≥ 0.3. Mice that did not fit the criteria were conditioned a second time. There was a minimum interval of 2 h between the retrieval and the FC HP test.

The FC HP test consisted of two trials, one in which the CS^+^ was presented and another in which the CS^-^ was presented. The two trials were counterbalanced within the group. The testing surface was set at 30 °C. After a 60 s acclimatization period, its temperature gradually increased at 6 °C per minute (HP start). Tone presentation (CS^+^ or CS^-^) started 130 s after the HP start. Temperature increase and tone presentation terminated concomitantly with the display of a nociception response. Valid nociception readout responses included jumping or licking the hindpaw. The effect of the negative emotional modulation on pain sensitivity was assessed by comparing the latency (or temperature) of the NC response between the two HP trials. There was a minimum of 30 min interval between the two trials for each mouse, during which mice returned to their home cage.

Freezing during the HP test was assessed by automatically analyzing thermographic camera videos using custom scripts (available at: https://github.com/djercog/TestoFreez). Briefly, individual video frames were binarized to define a mask for the mouse within each frame. Consecutive binarized frames were then subtracted, and the number of non-zero pixels in the resultant image determined if the mouse was considered immobile when below a certain threshold (defined to approximate manual freezing estimation on random videos). Immobilization episodes of at least 0.5 s were considered as freezing periods. In contrast, non-immobilization episodes shorter than 0.1 s were merged together with the neighbouring detected freezing episodes.

#### Extinction training

On days 3 and 4, mice (n =10) were submitted to an extinction training protocol established by Courtin et al. 2014^41^. Briefly, there were 4 CS^-^ and 12 CS^+^, presented in a non-reinforced manner (**Figure 1g**). Mice were considered to have successfully extinguished the fear expression if the level of freezing in the last 4 CS^+^ was not statistically different from the CS^-^. After fear extinction, mice were submitted to the FC HP test, using the same parameters as in the CFCA paradigm.

#### Stability training

The first three days consisted of the classical CFCA paradigm. On day 4, mice (n = 10) repeated the protocol applied on day 3 (**Figure 1—figure supplement 2e-i)**. All mice that met the fear conditioning criteria (DI ≥ 0.4 & CI ≥ 0.3) on day 3 were kept for the following day, independent of their level of Conditioning on day 4.

#### Vasoconstriction assay

To determine whether vasoconstriction was a confounding factor in the FC HP test outcomes, we used a modified version of the CFCA paradigm (**Figure 1f)**. During the HP test (n = 13), the temperature was not gradually increased but maintained at 30 °C while CSs were presented. The tones (CS^+^ or CS^-^) were presented for 120 s, and mice were kept for an additional 50 s before the HP trial terminated. Offline, with the infrared videos, mice’s back and tail were measured at 30 s intervals. For each measurement point, the temperatures of three spots on the back and tail were averaged to obtain the body temperature for each body part at that time point and mouse.

### Basal nociception assay (No tone)

Naïve mice (n = 10) were submitted to the HP test (**Figure 1i and Figure 1—figure supplement 2a-d)**. Each mouse underwent two identical trials. The HP test was similar to the one described in the CFCA paradigm, except no tones were presented. There was a minimum of 30 min interval between the two trials, during which mice returned to a restful state in their home cage. Since the two trials were not significantly different, for simplicity, in panel i of Figure 1, the two trials were pooled together.

#### Tone-specific nociception assay

Naïve mice (n = 12) were submitted to the HP test without auditory fear conditioning (**Figure 1i and Figure 1—figure supplement 2a-d)**. Each animal underwent two trials, one trial with a 7.5 kHz tone presentation and another with a WN tone presentation. The tone presentation was counterbalanced. The HP test parameters were similar to those described for the CFCA, except that both tones were unconditioned. Since the two trials were not significantly different, for simplicity, in panel i of Figure 1, the two trials were pooled together.

#### Open Field assay

This test was used to determine the effect of optogenetic stimulation on locomotion. Therefore, only mice from optogenetic experiments were submitted to this assay. Mice could freely move during the entire test. The locomotion assay had a total duration of 9 min and was divided into 3 min epochs. The first and third epochs were OFF periods in which no optical stimulation occurred. During the second epoch, mice received photostimulation (ON period). The optogenetic stimulation effect was analyzed by comparing the overall distance traveled between the OFF and ON epochs (**Figure 2—figure supplement 3c, d)**. Mice injected with GFP were used to test the effect of heat and light on the stimulation itself. Mouse tracking data were extracted from video recordings using idTracker^42^ and imported into Matlab for further analysis with custom scripts to calculate the total distance traveled during stimulation.

#### Real-time Place Preference assay

Only mice from optogenetic experiments were submitted to this assay. During the entire duration of the assay, mice could freely shuttle between the two compartments. Under closed-loop stimulation, mice received photo-stimulation upon entry into one of the two compartments. The stimulated compartment was counterbalanced within the group. The optical stimulation effect was assessed by comparing the time spent in the stimulated compartment between the ChR2 and GFP groups (**Figure 2—figure supplement 3e, f**). Permanence time in the stimulated compartment was automatically assessed by the apparatus software (Imetronic).

### Virus injections and optogenetics

For optogenetic manipulation of SST^+^ cells in the vlPAG, 0.15-0.2 μL of either ChR2 (AAV5-EF1a-DIO-hChR2(H134R)-EYFP, titer: 3.2×10^12^ - Vector Core, University of North Carolina), ArchT (AAV9-CAG-FLEX-ArchT-GFP, titer: 4.7×10^12^ - Vector Core, University of North Carolina) or GFP (AAV5-FLEX-GFP, titer: 4.5×10^12^ - Vector Core, University of North Carolina) were bilaterally injected into the vlPAG of 8/9 weeks old *Sst-IRES-Cre* mice from glass pipettes (tip diameter 20-30 μm) at the following coordinates relative to bregma: - 4.4 mm AP; ± 1.5 mm ML; −2.45 mm DV from dura, with a 20 degrees angle. Injection coordinates for manipulating VIP^+^ vlPAG cells were as follows: - 4.4 mm AP; ± 1.35 mm ML; −2.5 mm DV from dura, with a 20 degrees angle.

At two weeks after the injections, mice were bilaterally implanted with custom-built optic fibers (diameter: 200 µm; numerical aperture: 0.39; Thorlabs) above the vlPAG at the following coordinates relative to bregma: i) *Sst-IRES-Cre* mice: - 4.4 mm AP; ± 1.0 mm ML; −1.8 mm DV from dura, with a 10 degrees angle; ii) *Vip-IRES-Cre* mice: - 4.4 mm AP; ± 0.8 mm ML; −2.0 mm DV from dura, with a 10 degrees angle. For RVM manipulations, mice were implanted at the following coordinates relative to bregma: −5.8 mm AP; 0.0 mm ML; −5.2 DV from the dura. All implants were secured using three stainless steel screws and Super-Bond cement (Sun Medical). During surgery, long- and short-lasting analgesic agents were injected (Metacam, Boehringer; Lurocaïne, Vetoquinol). After surgery, mice were allowed to recover for at least five days. Afterward, mice were handled daily to familiarize themselves with being restrained for the optic fibers connection. Behavioral experiments were performed at least four weeks after viral injections. Only mice with correct placement of optic fibers and virus expression were included in the analyses.

For optogenetic excitation, light stimulation consisted of blue light (473 nm, ∼8-10 mW at fiber tip) delivered with 2 Hz frequency and 5 ms pulse duration. In contrast, light stimulation consisted of green light (532 nm, ∼8-10 mW at fiber tip) delivered continuously for optogenetic inhibition. For optogenetic manipulations during the CFCA paradigm of either SST^+^ or VIP^+^ vlPAG cells, the light was delivered during the FC HP test and paired with the tone presentation.

Two sets of experiments were performed sequentially for stimulation during the fear conditioning. First, the footshock US was replaced by optical stimulation. Then, the US became the optical stimulation combined with the footshock (**Figure 2—figure supplement 4**). For both conditions, optogenetic stimulation began 5 s before and lasted until 5 s after CS^+^ offset. The same mice were used for both experiments. For the manipulations during the fear retrieval, there were 12 CS^+^ presentations divided into blocks of 4 CS^+^. The optogenetic stimulation was paired with the second block of CS^+^.

### Single-molecule *in situ* hybridization

Expression of *Sst*, *Slc32a1*, *Slc17a6,* and *Cre* transcripts was detected in C57/BL6J mice using single-molecule fluorescent *in situ* hybridization (smFISH) as previously described (Castell et al., 2024 PMID: 37858736). Briefly, coronal PAG sections (16 µm) were collected directly onto Superfrost Plus slides. Probes for *Sst* (ACDBio; Cat#404631), *Slc32a1* (ACDBio; Cat#319191-C2), *Slc17a6* (ACDBio; Cat#319171-C2), and *Cre* (ACDBio; Cat#312281-C2) were used with the RNAscope Fluorescent Multiplex Kit (ACDBio; Cat# 323110). Fluorescence was captured using sequential laser scanning confocal microscopy (Leica SP8), and quantification was performed in ImageJ.

### Anatomical tracing

#### Fluorogold injection

An incision between one to two cm was made slightly caudal to the peak of the dorsal hump to expose the lumbar spinal region. The vertebra of interest was identified. Then, a small incision was made between the tendons and the vertebral column on either side. The vertebra was then secured using spinal adaptor clamps, and all tissue was removed from the surface of the bone. Pulled borosilicate glass capillaries (Ringcaps, disposable capillary pipettes with ring mark, DURAN, Hirschmann Laborgeräte, Germany) were inserted between 2 vertebrae and allowed to microinject 0.5 μL of fluorogold 2% in the dorsal horn of the spinal cord on both sides.

### *In vivo* electrophysiology

Mice were anesthetized with isoflurane 4% for induction, then 1.5% maintenance. The experiment was started as soon as there was no longer any reflex. The colorectal temperature was kept at 37 °C with a heating blanket. Two metal clamps were used to stabilize the animal’s spine in a stereotactic frame (M2E, France) during electrophysiological recordings. Then, a laminectomy was performed at T13-L1 to expose the lumbar part of the spinal cord. The dura mater was carefully removed. Custom-made optical fibers were placed 1mm above the dorsal spinal cord for optogenetic manipulations. C-fiber-evoked field potentials were recorded in the deep lamina of the DH (at a depth range of 250 and 500 µm) with borosilicate glass capillaries (2 MΩ, filled with NaCl 684 mM; Harvard Apparatus, Cambridge, MA, USA). Field potentials were recorded with an ISODAM-amplifier (low filter: 0.1Hz to high filter: 0.1 kHz; World Precision Instruments, USA) in response to electrical stimulation of the ipsilateral paw. Single unit recordings of WDR DH cells were made with the same borosilicate glass capillaries mentioned above and placed in the dorsal part of the spinal cord. The criterion for selecting a cell was the presence of an A-fiber-evoked response (0-80 ms) followed by a C-fiber-evoked response (80-150 ms) to electrical stimulation of the ipsilateral sciatic nerve.

The paw of the mice was stimulated by trains (every 30 s) of electrical stimulation at two times the threshold for C-fibers, which were performed before, during, and after optogenetic stimulations with an optic fiber placed above the recording site. Subthreshold stimulations were performed below the threshold for C-fiber and A-fiber, respectively.

### Histology analyses

Mice were administered a lethal dose of Exagon and underwent transcardial perfusions via the left ventricle with 4% w/v paraformaldehyde (PFA) in 0.1 M PB. Following dissection, brains were post-fixed for 24 h at 4°C in 4% PFA. Brain sections of 80 μm-thick were cut on a vibratome, mounted on gelatin-coated microscope slides, and dried. To verify correct viral injections and optic fiber location, serial 80 μm-thick slices containing the regions of interest were mounted in VectaShield (Vector Laboratories) and imaged using an epifluorescence system (Leica DM 5000) fitted with a 10-x dry objective. The location and the extent of the injections/infections were visually controlled. Only infections targeting the vlPAG and optic fibers terminating, depending on the experiment, above the vlPAG or RVM were included in the analyses.

### Statistics

All statistical details are contained in Supplementary File 1. Box-whisker plots indicate median, interquartile range, and 5th - 95th percentiles of the distribution. All statistics are indicated where used. Statistical analyses were performed with Prism software (GraphPad). We tested the normality of all data with the Kolmogorov–Smirnov normality test in Prism software (GraphPad), and used nonparametric statistical tests for nonnormally distributed datasets, as indicated where used. No statistical methods were used to predetermine sample sizes, but sample sizes were based on previous studies in our laboratory. Blinding for the opsin was done for optogenetics experiments. Animals used for the CFCA paradigm that did not satisfy the DI and CI criteria after two conditioning sessions were discarded from the study. Mice with an incorrect injection of the opsins or misplaced location of the optic fibers were discarded. For *in vivo* electrophysiology, field potentials were measured as the area above the curve in the C-fiber range (80-300 ms). Absolute values were used to compare values in each condition (OFF vs. ON optogenetic manipulation). In single-unit recordings, the number of A- and C-fiber induced spikes in WDR cells was measured after each electrical stimulation, during, and after optogenetic manipulation. An average of the four stimuli for each WDR recorded was used for statistical analysis. Outliers were removed using the Tukey method.

No other mice were excluded. Significance levels are indicated as follows: *p<0.05, **p<0.01, ***p<0.001, **** p<0.0001.

## Supporting information

Supplementary figures and statistical table

## Data availability

All data generated or analysed during this study, including the source data underlying every figure, have been deposited in the Dryad Digital Repository under DOI https://doi.org/10.5061/dryad.63xsj3vj3. Custom code used for the automated detection of freezing behaviour from thermographic recordings (TestoFreez)^43^ is available at https://github.com/djercog/TestoFreez, released under the MIT licence.

## Acknowledgments

We thank Joshua Johansen, David Finn, and Nadine Gogolla for critical and stimulating discussions; K. Deisseroth and E. Boyden for generously sharing material, S. Laumond, J. Tessaire, and the technical staff of the housing and experimental animal facility of the Neurocentre Magendie. We thank Claire Francioni for her technical support. Microscopy was performed in the Bordeaux Imaging Center of the CNRS-INSERM and Bordeaux University, member of France BioImaging. This work was supported by grants from the French National Research Agency (ANR-FEARLESSPAIN, ANR-10-EQPX-08 OPTOPATH), the Fondation pour la Recherche Médicale (FRM-PhD grant to NW 2020-2021) and the French Society for the Study and Treatment of Pain (SFETD – grant to NW 2022).

## Figure supplements

**Figure 1—figure supplement 1. Cued fear-conditioned analgesia behavior. a.** During habituation, the freezing levels for the context (BL) and the two tones were low but significantly different (***, P = 0.0003, Friedman test, n = 12 mice). **b.** Mean temperature at which the NC response was observed in the HP test. The temperature of NC response was higher during CS^+^ trials when compared to the CS^-^trials (***, P = 0.0005, Wilcoxon matched-pairs signed rank test, n = 12 mice)**. c.** Schematic representation of how the back and tail temperature of the mice was measured. Every 30-sec three-point temperature was taken for each of the two body parts. The temperature of the mice back and tail were measured by the infrared digital thermographic camera and analyzed offline (see Methods). The average temperature of the mice back (**d**) and tail (**e**) while the CS^+^ or the CS^-^were presented. There were no differences in body temperature for the different CSs trials (ns, P > 0.05, mixed-effects model, n = 13 mice). Vertical dashed lines correspond to the average time of NC response for the CS^+^ (red) and CS^-^(blue) during the standard CFCA protocol. Figure 1**—figure supplement 21. CFCA depends on associative processes. a. Top**, protocol for the *No Tone*: naïve mice were submitted to two HP trials without conditioning nor tone presentation. **Bottom**, protocol for the *Tone*: naïve mice were submitted to two HP trials paired with an unconditioned tone presentation. **b, c.** Mean latency and temperature of NC response for the two tests mentioned above and the CFCA. The transient cued-fear induced analgesia (CS^+^) compared to the basal nociception, the tone, and the CS^-^trials (***, P = 0.0001, Kruskal-Wallis test). Trials between the same type of test (*No tone* or *Tone*) were not significantly different (ns, P > 0.05, Bonferroni post-hoc test, see Supplementary File 1 for all comparisons) **d.** After fear extinction, there was no difference in the mean temperature response between the two CSs (ns, P = 0.3276 paired t-test, n = 10 mice). **e.** Protocol for stability training (see methods). Mice were submitted to two rounds of the CFCA paradigm. During retrieval (**f, g**), average freezing values during CS^+^ was higher than CS^-^ or baseline (BL) periods (****, P < 0.0001, one-way RM ANOVA, n = 10 mice). **h, i.** Mean latency and temperature of NC response during CS^-^ and CS^+^ trials for FC HP test 1 (**h**) and test 2 (**i**). CS^+^ presentation increased the latency and temperature of NC response in FC HP test 1 and test 2 (*, **, and ***, P < 0.05, paired t-test; for details, see Supplementary File 1).

**Figure 1—figure supplement 3. Two distinct neuronal populations in *Sst-IRES-Cre* mice within the PAG. a.** Representative picture of single-molecular fluorescent in situ hybridization for *Sst* mRNAs in the PAG. Scale bar, 400 µm. **b, c.** Single-molecular fluorescent in situ hybridization for *Sst* (green) and *Cre* (red) mRNAs in the vlPAG (left). Histograms showing the co-expression of *Sst/Cre* as percentage of *Sst*-expressing cells (green) and as percentage of *Cre*-expressing cells (red) in the dlPAG and vlPAG (right). Scale bar, 20 µm. In the dlPAG, 94% of RNA *Sst*-expressing-cells (101 out of 107 cells) expressed *Cre* RNA, and 84 % of RNA *Cre*-expressing cells expressed somatostatin RNA (101 out of 120 cells). In the vlPAG, the values are 88% (74 out of 84 cells) and 99% (74 out of 75 cells) for *Cre* and *Sst*, respectively. **d.** Single-molecular fluorescent in situ hybridization for *Sst* (red) and *Slc32a1* (green) within the dlPAG (upper panel) and vlPAG (bottom panel). Scale bar, 20 µm. **e.** Quantification of colocalization within the dlPAG (upper panel) and vlPAG (bottom panel) of *Sst*^+^ and *Slc32a1*^+^. In the dlPAG, approximately 21% of Slc32a1+ cells (17 out of 80 cells) are *Sst*^+^, and 19% (17 out of 88 cells) of *Sst*^+^ cells are *Slc32a1*^+^. On the contrary, in the vlPAG, approximately 98% of Slc32a1+ cells (183 out of 186 cells) are *Sst*^+^, and 72% of Sst+ cells (183 out of 254 cells) are *Slc32a1*^+^. **f.** Single-molecular fluorescent in situ hybridization for *Sst* (red) and *Slc17a6* (green) within the dlPAG (upper panel) and vlPAG (bottom panel). Scale bar, 20 µm. **g.** Quantification of colocalization within the dlPAG (upper panel) and vlPAG (bottom panel) of *Sst*^+^ and *Slc17a6*^+^. In the dlPAG, approximately 89% of Slc17a6+ cells (54 out of 61 cells) are *Sst*^+^, and 31% of Sst+ cells (54 out of 172 cells) are *Slc17a6*^+^. On the contrary, in the vlPAG, approximately 24% of Slc17a6+ cells (43 out of 176 cells) are *Sst*^+^ and 37% of Sst+ cells (43 out of 115 cells) are *Slc17a6*^+^. White arrows indicate colocalization.

**Figure 2—figure supplement 1. Optical fiber placement for *Sst-IRES-Cre* mice.** Anatomical location of the tip of the optic fiber for *Sst-IRES-Cre* mice (n = 39).

**Figure 2—figure supplement 2. Comparable fear levels prior to the FC HP test**. After retrieval, fear conditioning levels between the opsins and their respective GFP groups were tested to ensure equivalent fear levels. The discrimination index (see methods) between the GFP and ChR2 (**a**, ns, P = 0.6052 unpaired t-test, n = 16 mice) or ArchT (**b,** ns, P = 0.4575, unpaired t-test, n = 23 mice) were not significantly different. All the mice discriminated equally between CSs. The conditioning index (see methods) between the GFP and ChR2 (**c,** ns, P = 0.2333, unpaired t-test n = 16 mice) or ArchT (**d,** ns, P = 0.1320, unpaired t-test, n = 23 mice) were also not significantly different. All mice displayed a similar high freezing level to the CS^+^.

**Figure 2—figure supplement 3. Optogenetic manipulation of SST^+^ vlPAG cells does not affect locomotion or produce aversion. a.** Photoactivation of SST^+^ vlPAG cells abolished the analgesic effect induced during defensive states (**, P = 0.0045, opsin x CS - two-way RM ANOVA, n = 7 GFP, n = 9 ChR2). The mean temperature of NC response for the CS^+^ was significantly different between the ChR2 and GFP groups (*, P = 0.0322, Bonferroni post-hoc test). For the ChR2 group, the mean temperature of NC response during CS^+^ was equivalent to the CS^-^(ns, P = 0.1783, Bonferroni post-hoc test). On the contrary, the mean temperature of NC response between the CSs differed for the GFP group (****, P < 0.0001, Bonferroni post-hoc test). **b.** Photoinhibition of SST^+^ vlPAG cells increased the analgesic effect for the ArchT group when compared to the GFP (**, P = 0.0035, opsin x CS - two-way RM ANOVA, n = 12 GFP, n = 11 ArchT). The mean temperature of NC response for the CS^-^ was significantly different between the ArchT and GFP groups (****, P < 0.0001, Bonferroni post-hoc test). For the GFP group, the mean temperature of NC response was higher for the CS^+^ trials when compared to the CS^-^trials (***, P = 0.0003, Bonferroni post-hoc test), yet this was not the case for the ArchT group (ns, P > 0.999, Bonferroni post-hoc test). **c.** The photoactivation of SST^+^ vlPAG cells was performed during the ON epoch (blue shaded area), and the average distance traveled during the ON epoch was not different from the OFF epochs when comparing the two opsins (ns, P = 0.5121, opsin x light - two-way RM ANOVA, n = 7 GFP, n = 9 ChR2). **d.** The photoinhibition of SST^+^ vlPAG cells was performed during the ON epoch (yellow shaded area), and the average distance traveled during the ON epoch was not different from the OFF epochs when comparing the two opsins (ns, P =0.4047, opsin x light - two-way RM ANOVA, n = 12 GFP, n = 11 ArchT). **e.** Real-time place-preference location plot from a representative animal while submitted to optogenetic activation of SST^+^ vlPAG cells in the left compartment throughout the 15-min session. **f**. There was no difference between ChR2-expressing mice and the control group in the time spent in the stimulated and non-stimulated compartments (ns, P = 0.1812, Mann-Whitney test).

**Figure 2—figure supplement 4. SST^+^ vlPAG cells photoactivation during conditioning does not impact fear learning. a**. Protocol for optogenetic manipulations during fear conditioning. On Day 2, mice received 5 CS-US associations. The US was either optogenetic stimulation alone or optogenetic stimulation plus footshock. **b.** Average freezing levels during the CS-US association of optogenetic stimulation alone. There was no difference in the overall freezing levels for the CS^-^ and CS^+^ between ChR2 and GFP (ns, P = 0.4804, opsin x CS - two-way RM ANOVA, n = 6 GFP, n = 7 ChR2). **c.** During retrieval, there was no effect on freezing upon activation of SST^+^ vlPAG cells as an US (ns, P = 0.1695, opsin x CS effect - two-way RM ANOVA, n = 6 GFP, n = 7 ChR2). **d.** Average freezing levels during the CS-US association of optogenetic stimulation plus footshock. There was no difference in the overall freezing levels for the CS^+^ and CS^-^between ChR2 and GFP (ns, P = 0.8663, opsin x CS effect - two-way RM ANOVA, n = 6 GFP, n = 7 ChR2). **e.** During retrieval, there was no difference in the fear expression by the activation of the SST^+^ vlPAG cells (ns, P = 0.8967, opsin x CS effect - two-way RM ANOVA, n = 6 GFP, n = 7 ChR2).

**Figure 2—figure supplement 5. Optogenetic effect observed during CFCA is not due to alteration of somatostatin levels nor mediated by VIP^+^ vlPAG cells. a.** *Sst-IRES-Cre* heterozygotic mice received a bilateral injection of opsins in the vlPAG, and optic fibers were implanted above the vlPAG. **b.** A representative example of expression patterns of ChR2 within SST^+^ vlPAG cells. **c.** Average freezing values during retrieval. ChR2 and GFP groups were pooled together because no differences were found in the conditioning level (data not shown). The average freezing values during CS^+^ were higher than CS^-^ or baseline (BL) periods (****, P < 0.0001, Friedman test, n = 13 mice). **d.** Optogenetic activation of SST^+^ vlPAG cells abolished the analgesic effect induced by defensive states (**, P = 0.0048, opsin x CS - two-way RM ANOVA, n = 6 GFP, n = 7 ChR2). The mean latency of NC response for the CS^+^ was significantly different between the ChR2 and GFP groups (*, P = 0.0300, Bonferroni post-hoc test, n = 6 GFP, n = 7 ChR2). For the ChR2 group, the latency of NC response during CS^+^ was equivalent to the CS^-^(ns, P = 0.3342, Bonferroni post-hoc test). On the contrary, the latency of NC response between the CSs was different for the GFP group (****, P = 0.0001, Bonferroni post-hoc test). **e.** *Vip-IRES-Cre* mice received a bilateral injection of opsins in the vlPAG, and optic fibers were implanted above the vlPAG. **f.** Representative example of the expression pattern of ArchT within VIP^+^ vlPAG cells. **g.** ArchT and GFP groups were pooled together because no differences were found in the conditioning level (data not shown). The average freezing values during CS^+^ were higher than CS^-^ or BL periods (****, P < 0.0001, one-way RM ANOVA, n = 10 mice). **h.** Optogenetic inhibition of VIP^+^ cells did not change the analgesic effect for the ArchT group when compared to the GFP (ns, P = 0.5455, opsin x CS effect - two-way RM ANOVA, n = 5 GFP, n = 5 ArchT). The latency of NC response was higher for the CS^+^ trials compared to the CS^-^trials for both GFP and ArchT (***, P < 0.01, Bonferroni post-hoc test, n = 5 GFP, n = 5 ArchT).

**Figure 2—figure supplement 6. Optical fiber placement for *Vip-IRES-Cre*.** Anatomical location of the tip of the optic fiber for *Vip-IRES-Cre* mice (n = 10).

**Figure 4—figure supplement 1. Optogenetic activation of SST^+^ vlPAG cells did not affect tactile sensitivity. a.** Perievent stimulus time histogram indicating the spiking latency of a purely tactile neuron recorded in the spinal cord in response to gentle brushing of the skin. **b.** Representative spiking activity latency of a purely tactile neuron recorded in the spinal cord in response to gentle brushing of the skin with or without activation of ChR2 expressing SST^+^ vlPAG cells. **c.** The activation of ChR2 expressing SST^+^ vlPAG cells of tactile neurons recorded in the spinal cord in response to gentle brushing of the skin had no effect on the average firing activity (Paired t-test, P = 0.6929, n = 5 cells).

**Figure 5—figure supplement 1. Optogenetic manipulation of GFP expressing vlPAG cells did not affect analgesia. a.** Single-unit recordings of WDR cells in the lumbar spinal cord during optogenetic manipulation of GFP expressing SST^+^ vlPAG inputs to the RVM. **b**. Photostimulation of SST^+^ vlPAG inputs to the RVM has no effect.

## References

1. Bolles, R. C. & Fanselow, M. S. A perceptual-defensive-recuperative model of fear and pain. Behav. Brain Sci. 3, 291–301 (1980).

2. Rhudy, J. L. & Meagher, M. W. Fear and anxiety: divergent effects on human pain thresholds. Pain 84, 65–75 (2000).

3. Flor, H. & Grösser, S. M. Conditioned stress-induced analgesia in humans. Eur. J. Pain 3, 317–324 (1999).

4. Beckham, J. C. et al. Chronic posttraumatic stress disorder and chronic pain in Vietnam combat veterans. J. Psychosom. Res. 43, 379–389 (1997).

5. International Association for the Study of Pain. IASP Terminology. Iasp (2017).

6. Butler, R. K. & Finn, D. P. Stress-induced analgesia. Prog. Neurobiol. 88, 184–202 (2009).

7. Rea, K., Roche, M. & Finn, D. P. Modulation of Conditioned Fear, Fear-Conditioned Analgesia, and Brain Regional C-Fos Expression Following Administration of Muscimol into the Rat Basolateral Amygdala. J. Pain 12, 712–721 (2011).

8. Floyd, N. S., Price, J. L., Ferry, A. T., Keay, K. A. & Bandler, R. Orbitomedial prefrontal cortical projections to distinct longitudinal columns of the periaqueductal gray in the rat. J. Comp. Neurol. 422, 556–578 (2000).

9. LeDoux, J., Iwata, J., Cicchetti, P. & Reis, D. Different projections of the central amygdaloid nucleus mediate autonomic and behavioral correlates of conditioned fear. J. Neurosci. 8, 2517–2529 (1988).

10. Vertes, R. P. Differential projections of the infralimbic and prelimbic cortex in the rat. Synapse 51, 32–58 (2004).

11. Carrive, P. & Morgan, M. M. Periaqueductal Gray - Chapter 10. The Human Nervous 631 System (Elsevier, 2012). doi:10.1016/B978-0-12-374236-0.10010-0.

12. Basbaum, A. I. & Fields, H. L. Endogenous Pain Control Systems: Brainstem Spinal Pathways and Endorphin Circuitry. Annu. Rev. Neurosci. 7, 309–338 (1984).

13. Millan, M. J. Descending control of pain. Prog. Neurobiol. 66, 355–474 (2002).

14. Fields, H. L., Heinricher, M. M. & Mason, P. Neurotransmitters in Nociceptive Modulatory Circuits. Annu. Rev. Neurosci. 14, 219–245 (1991).

15. Samineni, V. K. et al. Divergent Modulation of Nociception by Glutamatergic and GABAergic Neuronal Subpopulations in the Periaqueductal Gray. eneuro 4, ENEURO.0129-16.2017 (2017).

16. V., C., C., M. & M., B. Role of benzodiazepine and serotonergic mechanisms in conditioned freezing and antinociception using electrical stimulation of the dorsal periaqueductal gray as unconditioned stimulus in rats. Psychopharmacology (Berl.) 165, 77–85 (2002).

17. Helmstetter, F. J. & Landeira-Fernandez, J. Conditional hypoalgesia is attenuated by Naltrexone applied to the periaqueductal gray. Brain Res. 537, 88–92 (1990).

18. Bellgowan, P. S. F. & Helmstetter, F. J. The role of mu and kappa opioid receptors within the periaqueductal gray in the expression of conditional hypoalgesia. Brain Res. 791, 83–89 (1998).

19. Jacquet, Y. F. The NMDA receptor: central role in pain inhibition in rat periaqueductal gray. Eur. J. Pharmacol. 154, 271–276 (1988).

20. Moreau, J. L. & Fields, H. L. Evidence for GABA involvement in midbrain control of medullary neurons that modulate nociceptive transmission. Brain Res. 397, 37–46 (1986).

21. Osborne, P. B., Vaughan, C. W., Wilson, H. I. & Christie, M. J. Opioid inhibition of rat periaqueductal grey neurones with identified projections to rostral ventromedial medulla in vitro. J. Physiol. 490, 383–389 (1996).

22. Harris, J. A. Descending antinociceptive mechanisms in the brainstem: Their role in the animal’s defensive system. J. Physiol.-Paris 90, 15–25 (1996).

23. Tovote, P. et al. Midbrain circuits for defensive behaviour. Nature 534, 206–212 (2016).

24. Lau, B. K. & Vaughan, C. W. Descending modulation of pain: the GABA disinhibition hypothesis of analgesia. Curr. Opin. Neurobiol. 29, 159–164 (2014).

25. Fields, H. State-dependent opioid control of pain. Nat. Rev. Neurosci. 5, 565–575 (2004).

26. Ossipov, M. H., Dussor, G. O. & Porreca, F. Central modulation of pain. J. Clin. Invest. 120, 3779–3787 (2010).

27. Vianna, D. M. L. & Carrive, P. Changes in cutaneous and body temperature during and after conditioned fear to context in the rat. Eur. J. Neurosci. 21, 2505–2512 (2005).

28. Smith, G. S. T. et al. Distribution of messenger RNAs encoding enkephalin, substance P, somatostatin, galanin, vasoactive intestinal polypeptide, neuropeptide Y, and calcitonin gene-related peptide in the midbrain periaqueductal grey in the rat. J. Comp. Neurol. 350, 23–40 (1994).

29. Viollet, C. et al. Somatostatin-IRES-Cre Mice: Between Knockout and Wild-Type? Front. Endocrinol. 8, 131 (2017).

30. Gasser, H. S. The classification of nerve fibers. Ohio J. Sci. (1941).

31. Mogil, J. S. Sex differences in pain and pain inhibition: multiple explanations of a controversial phenomenon. Nat. Rev. Neurosci. 13, 859–866 (2012).

32. Aby, F. et al. Switch of serotonergic descending inhibition into facilitation by a spinal chloride imbalance in neuropathic pain. Sci. Adv. 8, eabo0689 (2022).

33. Grivet, Z. et al. Brainstem serotonin amplifies nociceptive transmission in a mouse model of Parkinson’s disease. Npj Park. Dis. 11, 11 (2025).

34. Tovote, P., Fadok, J. P. & Lüthi, A. Neuronal circuits for fear and anxiety. Nat. Rev. Neurosci. 16, 317–331 (2015).

35. Yin, W. et al. A Central Amygdala–Ventrolateral Periaqueductal Gray Matter Pathway for Pain in a Mouse Model of Depression-like Behavior. Anesthesiology 132, 1175–1196 (2020).

36. Zhang, Y. et al. Somatostatin Neurons from Periaqueductal Gray to Medulla Facilitate Neuropathic Pain in Male Mice. J. Pain 24, 1020–1029 (2023).

37. Morgan, M. M., Whittier, K. L., Hegarty, D. M. & Aicher, S. A. Periaqueductal gray neurons project to spinally projecting GABAergic neurons in the rostral ventromedial medulla. Pain 140, 376–386 (2008).

38. Butler, R. K. et al. Molecular and electrophysiological changes in the prefrontal cortex–amygdala–dorsal periaqueductal grey pathway during persistent pain state and fear-conditioned analgesia. Physiol. Behav. 104, 1075–1081 (2011).

39. Tovote, P., Esposito, M. S., Botta, P., Chaudun, F. & Jonathan, P. Midbrain circuits for defensive behaviour. Nat. Press https://doi.org/10.1038/nature17996 (2016) doi:10.1038/nature17996.

40. McWilliams, L. A., Cox, B. J. & Enns, M. W. Mood and anxiety disorders associated with chronic pain: an examination in a nationally representative sample. Pain 106, 127–133 (2003).

41. Courtin, J. et al. Prefrontal parvalbumin interneurons shape neuronal activity to drive fear expression. Nature 505, 92–6 (2014).

42. Pérez-Escudero, A., Vicente-Page, J., Hinz, R. C., Arganda, S. & De Polavieja, G. G. idTracker: tracking individuals in a group by automatic identification of unmarked animals. Nat. Methods 11, 743–748 (2014).

43. Jercog, D. TestoFreez: mouse freezing behavior detection from thermographic camera videos. GitHub (2026). https://github.com/djercog/TestoFreez

