## Supplementary figures and statistical table for "Midbrain somatostatin-expressing cells control pain-suppression during defensive states"

### Supplementary File 1 | Summary of statistical results associated with the Figures and figure supplements

| Figure number | Normality test | Statistic method | Main effect | Post hoc multiple comparisons test | Number of mice | Number of cells |
|---|---|---|---|---|---|---|
| 1b | Passed | One-way RM ANOVA followed by Bonferroni post-hoc test | $F_{(2, 22)}$ = 160.8<br>P value < 0.0001 | BL vs. $CS^-$, P > 0.9999, ns<br>BL vs. $CS^+$, P < 0.0001<br>$CS^-$ vs. $CS^+$, P < 0.0001 | 12 | NA |
| 1e | Failed | Wilcoxon matched-pairs signed rank test (two-tailed) | P = 0.0005 | *W* = 78.00 | 12 | NA |
| 1f | Onset, failed<br>Offset, failed | Wilcoxon matched-pairs signed rank test (two-tailed) | Onset, P = 0.0015<br>Offset, P = 0.8330, ns | Onset, *W* = 74<br>Offset, *W* = 6 | 12 | NA |
| 1g | Failed | Friedman test followed by Dunn's multiple post-hoc test | P value < 0.0001<br>*Q* = 48.13 | BL vs. $CS^-$, P > 0.9999, ns<br>BL/$CS^-$ vs. $CS^+_{1st\ block}$, P < 0.0001<br>$CS^-$ vs. $CS^+_{5th/6th\ block}$, P > 0.9999, ns<br>$CS^+_{1st\ block}$ vs. $CS^+_{5th/6th\ block}$, P < 0.005 | 10 | NA |
| 1h | Passed | Paired t-test (two-tailed) | P = 0.4365, ns | $t_9$ = 0.8143 | 10 | NA |
| 1i | Failed | Kruskal-Wallis test followed by Dunn's multiple post-hoc test | P < 0.0001<br>*H* = 25.32 | No tone vs. Tone, P = 0.7710, ns<br>No tone vs. $CS^-$, P > 0.9999, ns<br>Tone vs. $CS^-$, P > 0.9999, ns<br>No tone vs. $CS^+$, P < 0.0001<br>Tone vs. $CS^+$, P = 0.0009<br>$CS^-$ vs $CS^+$, P = 0.0033 | No tone = 10<br>Tone = 12<br>CSs = 12 | NA |
| 2c | Passed | One-way RM ANOVA followed by Bonferroni post-hoc test | $F_{(2,30)}$ = 396.8<br>P < 0.0001 | BL vs. $CS^-$, P > 0.9999, ns<br>BL vs. $CS^+$, P < 0.0001<br>$CS^-$ vs. $CS^+$, P < 0.0001 | 16 | NA |
| 2d | Passed | Two-way RM ANOVA followed by Bonferroni post-hoc test | Opsin x CS, $F_{(1,14)}$ = 8.514<br>P = 0.0112 | $CS^-$ vs. $CS^+$, GFP, P = 0.0003<br>$CS^-$ vs. $CS^+$, ChR2, P = 0.3876, ns<br>GFP vs. ChR2, $CS^-$, P > 0.9999, ns<br>GFP vs. ChR2, $CS^+$, P = 0.0344 | GFP = 7<br>ChR2 = 9 | NA |
| 2e | Passed | One-way RM ANOVA followed by Bonferroni post-hoc test | $F_{(2,44)}$ = 280.4<br>P < 0.0001 | BL vs. $CS^-$, P = 0.2337, ns<br>BL vs. $CS^+$, P < 0.0001<br>$CS^-$ vs. $CS^+$, P < 0.0001 | 23 | NA |
| 2f | Passed | Two-way RM ANOVA followed by Bonferroni post-hoc test | Opsin x CS, $F_{(1,21)}$ = 10.64<br>P = 0.0037 | $CS^-$ vs. $CS^+$, GFP, P = 0.0003<br>$CS^-$ vs. $CS^+$, ArchT, P > 0.9999, ns<br>GFP vs. ArchT, $CS^-$, P < 0.0001<br>GFP vs. ArchT, $CS^+$, P = 0.0265 | GFP = 12<br>ArchT = 11 | NA |
| 3c | Passed | Two-way RM ANOVA followed by Bonferroni post-hoc test | Opsin x CS, $F_{(4,48)}$ = 0.1954<br>P = 0.9396, ns | BL vs. $CS^-$, GFP/ArchT, P > 0.9999, ns<br>BL vs. $CS^+_{1st/2nd/3rd}$, GFP/ArchT, P < 0.0001<br>$CS^-$ vs. $CS^+_{1st/2nd/3rd}$, GFP/ArchT, P < 0.0001<br>$CS^+_{1st}$ vs. $CS^+_{2nd/3rd}$, GFP/ArchT, P > 0.9999, ns<br>$CS^+_{2nd}$ vs. $CS^+_{3rd}$, GFP/ArchT, P > 0.9999, ns | GFP = 6<br>ArchT = 8 | NA |

**NA = Not applicable NS = Not significant fs = Figure supplement**

### Supplementary File 1 | Summary of statistical results associated with the Figures and figure supplements

| Figure number | Normality test | Statistic method | Main effect | Post hoc multiple comparisons test | Number of mice | Number of cells |
|---|---|---|---|---|---|---|
| | | | | GFP vs. ArchT, BL/$CS^-$/$CS^+_{1st/2nd/3rd}$, P > 0.9999, ns | | |
| 3d | Passed | Two-way RM ANOVA followed by Bonferroni post-hoc test | Opsin x CS, $F_{(4,60)}$ = 7.957<br>P < 0.0001 | BL vs. $CS^-$, GFP/ChR2, P > 0.9999, ns<br>BL vs. $CS^+_{1st/2nd/3rd}$, GFP, P < 0.0001<br>$CS^-$ vs. $CS^+_{1st/2nd/3rd}$, GFP, P < 0.0001<br>$CS^+_{1st}$ vs. $CS^+_{2nd/3rd}$, GFP, P > 0.9999, ns<br>$CS^+_{2nd}$ vs. $CS^+_{3rd}$, GFP, P > 0.9999, ns<br>BL vs. $CS^+_{1st/3rd}$, ChR2, P < 0.0001<br>BL vs. $CS^+_{2nd}$, ChR2, P = 0.0789, ns<br>$CS^-$ vs. $CS^+_{1st/3rd}$, ChR2, P < 0.0001<br>$CS^-$ vs. $CS^+_{2nd}$, ChR2, P > 0.9999, ns<br>$CS^+_{1st}$ vs. $CS^+_{2nd}$, ChR2, P < 0.0001<br>$CS^+_{1st}$ vs. $CS^+_{3rd}$, ChR2, P = 0.2372, ns<br>$CS^+_{2nd}$ vs. $CS^+_{3rd}$, ChR2, P < 0.0001<br>GFP vs. ChR2, BL/$CS^-$/$CS^+_{1st/3rd}$, P > 0.9999, ns<br>GFP vs. ChR2, $CS^+_{2nd}$, P = 0.0002 | GFP = 9<br>ChR2 = 8 | NA |
| 4b | Passed | One-way RM ANOVA followed by Bonferroni post-hoc test | $F_{(2, 14)}$ = 17.94<br>P = 0.0001 | OFF1 vs. ON, P = 0.0006<br>OFF2 vs. ON, P = 0.0003<br>OFF1 vs. OFF2, P > 0.9999, ns | 8 | NA |
| 4c | Passed | One-way RM ANOVA followed by Bonferroni post-hoc test | $F_{(2, 8)}$ = 5.826<br>P = 0.0275 | OFF1 vs. ON, P = 0.0414<br>OFF2 vs. ON, P = 0.0452<br>OFF1 vs. OFF2, P = 0.9980, ns | 5 | NA |
| 4d | A-β fiber, passed<br>A-δ fiber, passed<br>C fiber, passed | Paired t-test (two-tailed) | A-β fiber, P = 0.5430, ns<br>A-δ fiber, P = 0.0022<br>C fiber, P = 0.0033 | A-β fiber, $t_{11}$ = 0.6319<br>A-δ fiber, $t_{11}$ = 3.959<br>C fiber, $t_{11}$ = 3.739 | 8 | A-β fiber, 12<br>A-δ fiber, 12<br>C fiber, 12 |
| 4e | A-β fiber, passed<br>A-δ fiber, passed<br>C fiber, passed | Paired t-test (two-tailed) | A-β fiber, P = 0.1386, ns<br>A-δ fiber, P = 0.0241<br>C fiber, P < 0.0001 | A-β fiber, $t_{13}$ = 1.578<br>A-δ fiber, $t_{13}$ = 2.553<br>C fiber, $t_{13}$ = 6.274 | 5 | A-β fiber, 14<br>A-δ fiber, 14<br>C fiber, 14 |
| 4f | A-β fiber, failed<br>A-δ fiber, failed<br>C fiber, failed | Wilcoxon matched-pairs signed rank test (two-tailed) | A-β fiber, P = 0.3750, ns<br>A-δ fiber, P = 0.0156<br>C fiber, P = 0.0020 | A-β fiber, $W$ = 7.0<br>A-δ fiber, $W$ = 28.0<br>C fiber, $W$ = 55.0 | 8 | A-β fiber, 10<br>A-δ fiber, 10<br>C fiber, 10 |
| 5e | Passed | Paired t-test (two-tailed) | P = 0.3683, ns | $t_{10}$ = 0.94222 | 7 | 11 |
| 5f, left | A-β fiber, passed<br>A-δ fiber, passed<br>C fiber, failed | Paired t-test (two-tailed)<br>Paired t-test (two-tailed)<br>Wilcoxon matched-pairs signed rank test (two-tailed) | A-β fiber, P = 0.0740, ns<br>A-δ fiber, P = 0.0346<br>C fiber, P = 0.0020 | A-β fiber, $t_9$ = 2.021<br>A-δ fiber, $t_9$ = 2.487<br>C fiber, $W$ = 55.0 | 7 | A-β fiber, 10<br>A-δ fiber, 10<br>C fiber, 10 |
| 5f, right | Failed | Wilcoxon matched-pairs signed rank test (two-tailed) | P = 0.0078 | $W$ = 36.0 | 7 | 8 |

**NA = Not applicable    NS = Not significant    fs = Figure supplement**

### Supplementary File 1 | Summary of statistical results associated with the Figures and figure supplements

| Figure number | Normality test | Statistic method | Main effect | Post hoc multiple comparisons test | Number of mice | Number of cells |
|---|---|---|---|---|---|---|
| 6b | Passed | Two-way RM ANOVA followed by Bonferroni post-hoc test | Opsin x CS, $F_{(4,44)} = 0.6292$<br>P = 0.6443, ns | BL vs. $CS^-$, GFP/ChR2, P > 0.9999, ns<br>BL vs. $CS^+_{1st/2nd/3rd}$, GFP/ChR2, P < 0.0001<br>$CS^-$ vs. $CS^+_{1st/2nd/3rd}$, GFP/ChR2, P < 0.0001<br>$CS^+_{1st}$ vs. $CS^+_{2nd}$, GFP/ChR2, P > 0.9999, ns<br>$CS^+_{2nd}$ vs. $CS^+_{3rd}$, GFP/ChR2, P > 0.9999, ns<br>$CS^+_{1st}$ vs. $CS^+_{3rd}$, GFP, P = 0.0090<br>$CS^+_{1st}$ vs. $CS^+_{3rd}$, ChR2, P = 0.0003<br>GFP vs. ChR2, BL/$CS^-$/$CS^+_{1st/2nd/3rd}$, P > 0.9999, ns | GFP = 7<br>ChR2 = 6 | NA |
| 6c | Passed | One-way RM ANOVA followed by Bonferroni post-hoc test | $F_{(2,26)} = 274.2$<br>P < 0.0001 | BL vs. $CS^-$, P > 0.9999, ns<br>BL vs. $CS^+$, P < 0.0001<br>$CS^-$ vs. $CS^+$, P < 0.0001 | 14 | NA |
| 6d | Passed | Two-way RM ANOVA followed by Bonferroni post-hoc test | Opsin x CS, F(1,12) = 19.87<br>P = 0.0008 | $CS^-$ vs. $CS^+$ - GFP, P < 0.0001<br>$CS^-$ vs. $CS^+$ - ChR2, P > 0.9999, ns<br>GFP vs. ChR2 - CS-, P = 0.1794, ns<br>GFP vs. ChR2 - CS+, P < 0.0001 | GFP = 7<br>ChR2 = 7 | NA |
| 1—fs1a | Failed | Friedman test followed by Dunn's multiple post-hoc test | P = 0.0003<br>*Q* = 16.55 | BL vs. $CS^-$, P = 0.3074, ns<br>BL vs. $CS^+$, P = 0.0003<br>$CS^-$ vs. $CS^+$, P = 0.0742, ns | 12 | NA |
| 1—fs1b | Failed | Wilcoxon matched-pairs signed rank test (two-tailed) | P = 0.0005 | *W* = 78.0 | 12 | NA |
| 1—fs1d | Passed | Mixed-effects model followed by Bonferroni post-hoc test | CS x Time, $F_{(9,238)} = 0.1716$<br>P = 0.9967 | $CS^+$ vs. $CS^-$ for each time point, all P > 0.9999, ns | 13 | NA |
| 1—fs1e | Passed | Mixed-effects model followed by Bonferroni post-hoc test | CS x Time, $F_{(9,238)} = 0.7147$<br>P = 0.6952 | $CS^+$ vs. $CS^-$ for each time point, all P > 0.9999, ns | 13 | NA |
| 1—fs2b | No tone, passed<br>Tone, passed<br>CFCA, failed | Paired t-test (two-tailed)<br>Paired t-test (two-tailed)<br>Wilcoxon matched-pairs signed rank test (two-tailed) | No tone, P = 0.9866, ns<br>Tone, P = 0.9085, ns<br>CFCA, P = 0.0005 | No tone, $t_9 = 0.01732$<br>Tone, $t_{11} = 0.1176$<br>CFCA, *W* = 78.0 | No tone = 10<br>Tone = 12<br>CFCA = 12 | NA |
| 1—fs2b | Failed | Kruskal-Wallis test followed by Dunn's multiple post-hoc test | P = 0.0001<br>*H* = 25.45 | No tone 1 vs. No tone 2/Tone 1/Tone 2/$CS^-$, P > 0.9999, ns<br>No tone 2 vs. Tone 1/Tone 2/$CS^-$, P > 0.9999, ns<br>Tone 1 vs. Tone 2/$CS^-$, P > 0.9999, ns<br>Tone 2 vs. Tone $CS^-$, P > 0.9999, ns<br>No tone 1 vs. $CS^+$, P = 0.0002<br>No tone 2 vs. $CS^+$, P = 0.0009<br>Tone 1 vs. $CS^+$, P = 0.0178<br>Tone 2 vs. $CS^+$, P = 0.0133<br>$CS^-$ vs. $CS^+$, P = 0.0082 | No tone = 10<br>Tone = 12<br>CFCA = 12 | NA |

**NA = Not applicable NS = Not significant fs = Figure supplement**

### Supplementary File 1 | Summary of statistical results associated with the Figures and figure supplements

| Figure number | Normality test | Statistic method | Main effect | Post hoc multiple comparisons test | Number of mice | Number of cells |
|---|---|---|---|---|---|---|
| 1—fs2c | No tone, passed<br>Tone, failed<br>CFCA, failed | Paired t-test (two-tailed)<br>Wilcoxon matched-pairs signed rank test (two-tailed) | No tone, P = 0.6216, ns<br>Tone, P = 0.4375, ns<br>CS, P = 0.0005 | No tone, $t_9$ = 0.5111<br>Tone, $W$ = -21.0<br>CFCA, $W$ = 78.0 | No tone = 10<br>Tone = 12<br>CFCA = 12 | NA |
| 1—fs2c | Failed | Kruskal-Wallis test followed by Dunn's multiple post-hoc test | P < 0.0001<br>$H$ = 29.30 | No tone 1 vs. No tone 2/Tone 1/Tone 2/$CS^-$, P > 0.9999, ns<br>No tone 2 vs. Tone 1/Tone 2/$CS^-$, P > 0.9999, ns<br>Tone 1 vs. Tone 2/$CS^-$, P > 0.9999, ns<br>Tone 2 vs. Tone $CS^-$, P > 0.9999, ns<br>No tone 1 vs. $CS^+$, P < 0.0001<br>No tone 2 vs. $CS^+$, P = 0.0010<br>Tone 1 vs. $CS^+$, P = 0.0339<br>Tone 2 vs. $CS^+$, P = 0.0037<br>$CS^-$ vs. $CS^+$, P = 0.0033 | No tone = 10<br>Tone = 12<br>CFCA = 12 | NA |
| 1—fs2d | Passed | Paired t-test (two-tailed) | P = 0.3276, ns | $t_9$ = 1.035 | 10 | NA |
| 1—fs2f | Passed | One-way RM ANOVA followed by Bonferroni post-hoc test | $F_{(2,18)}$ = 558.2<br>P < 0.0001 | BL vs. $CS^-$, P > 0.9999, ns<br>BL vs. $CS^+$, P < 0.0001<br>$CS^-$ vs. $CS^+$, P < 0.0001 | 10 | NA |
| 1—fs2g | Passed | One-way RM ANOVA followed by Bonferroni post-hoc test | $F_{(2,18)}$ = 56.00<br>P < 0.0001 | BL vs. $CS^-$, P > 0.9999, ns<br>BL vs. $CS^+$, P < 0.0001<br>$CS^-$ vs. $CS^+$, P < 0.0001 | 10 | NA |
| 1—fs2h, left | Passed | Paired t-test (two-tailed) | P = 0.0029 | $t_9$ = 4.054 | 10 | NA |
| 1—fs2h, right | Passed | Paired t-test (two-tailed) | P = 0.0006 | $t_9$ = 5.184 | 10 | NA |
| 1—fs2i, left | Passed | Paired t-test (two-tailed) | P = 0.0362 | $t_9$ = 2.459 | 10 | NA |
| 1—fs2i, right | Passed | Paired t-test (two-tailed) | P = 0.0245 | $t_9$ = 2.697 | 10 | NA |
| 1—fs3c | NA | Cell counts, no statistical test applied | dlPAG, ~94% (101/107) and ~84% (101/120)<br>vlPAG, ~88% (74/84) and ~99% (74/75) | NA | NA | dlPAG = 107 / 120<br>vlPAG = 84 / 75 |
| 1—fs3e | NA | Cell counts, no statistical test applied | dlPAG, ~21% (17/80) and ~19% (17/88)<br>vlPAG, ~98% (183/186) and ~72% (183/254) | NA | NA | dlPAG = 80 / 88<br>vlPAG = 186 / 254 |
| 1—fs3g | NA | Cell counts, no statistical test applied | dlPAG, ~89% (54/61) and ~31% | NA | NA | dlPAG = 61 / |

**NA = Not applicable NS = Not significant fs = Figure supplement**

### Supplementary File 1 | Summary of statistical results associated with the Figures and figure supplements

| Figure number | Normality test | Statistic method | Main effect | Post hoc multiple comparisons test | Number of mice | Number of cells |
|---|---|---|---|---|---|---|
| | | | (54/172)<br>vlPAG, ~24% (43/176) and ~37% (43/115) | | | 172<br>vlPAG = 176 / 115 |
| 2—fs1 | NA | No statistical test (anatomical verification of optic fiber placement) | NA | NA | 39 | NA |
| 2—fs2a | Passed | Unpaired t-test (two-tailed) | P = 0.6052 | $t_{14}$ = 0.5288 | GFP = 7<br>ChR2 = 9 | NA |
| 2—fs2b | Passed | Unpaired t-test (two-tailed) | P = 0.4575 | $t_{21}$ = 0.7569 | GFP = 12<br>ArchT = 11 | NA |
| 2—fs2c | Passed | Unpaired t-test (two-tailed) | P = 0.2333 | $t_{14}$ = 1.246 | GFP = 7<br>ChR2 = 9 | NA |
| 2—fs2d | Passed | Unpaired t-test (two-tailed) | P = 0.1320 | $t_{21}$ = 1.567 | GFP = 12<br>ArchT = 11 | NA |
| 2—fs3a | Passed | Two-way RM ANOVA followed by Bonferroni post-hoc test | Opsin x CS, $F_{(1,14)}$ = 11.40<br>P = 0.0045 | $CS^-$ vs. $CS^+$, GFP, P < 0.0001<br>$CS^-$ vs. $CS^+$, ChR2, P = 0.1783, ns<br>GFP vs. ChR2, $CS^-$, P > 0.9999, ns<br>GFP vs. ChR2, $CS^+$, P = 0.0322 | GFP = 7<br>ChR2 = 9 | NA |
| 2—fs3b | Passed | Two-way RM ANOVA followed by Bonferroni post-hoc test | Opsin x CS, $F_{(1,21)}$ = 10.84<br>P = 0.0035 | $CS^-$ vs. $CS^+$, GFP, P = 0.0003<br>$CS^-$ vs. $CS^+$, ArchT, P > 0.9999, ns<br>GFP vs. ArchT, $CS^-$, P < 0.0001<br>GFP vs. ArchT, $CS^+$, P = 0.0704, ns | GFP = 12<br>ArchT = 11 | NA |
| 2—fs3c | Passed | Two-way RM ANOVA followed by Bonferroni post-hoc test | Opsin x Light, $F_{(2,28)}$ = 0.6855<br>P = 0.5121, ns | OFF1 vs. ON, GFP/ChR2, P > 0.9999, ns<br>OFF2 vs. ON, GFP/ChR2, P > 0.9999, ns<br>OFF1 vs. OFF2, GFP/ChR2, P > 0.9999, ns<br>GFP vs. ChR2, OFF1/ON/OFF2, P > 0.9999, ns | GFP = 7<br>ChR2 = 9 | NA |
| 2—fs3d | Passed | Two-way RM ANOVA followed by Bonferroni post-hoc test | Opsin x Light, $F_{(2,42)}$ = 0.9243<br>P = 0.4047, ns | OFF1 vs. ON, GFP, P = 0.0165<br>OFF1 vs. ON, ArchT, P = 0.0003<br>OFF2 vs. ON, GFP/ArchT, P > 0.9999, ns<br>OFF1 vs. OFF2, GFP/ArchT, P > 0.9999, ns<br>GFP vs. ArchT, OFF1/ON/OFF2, P > 0.9999, ns | GFP = 12<br>ArchT = 11 | NA |
| 2—fs3f | Failed | Mann-Whitney test (two-tailed) | P = 0.1812 | $U$ = 13 | GFP = 6<br>ChR2 = 8 | NA |
| 2—fs4b | Passed | Two-way RM ANOVA followed by Bonferroni post-hoc test | Opsin x CS, $F_{(2,22)}$ = 0.7582<br>P = 0.4804, ns | GFP vs. ChR2, BL/$CS^-$/$CS^+$, P > 0.9999, ns | GFP = 6<br>ChR2 = 7 | NA |
| 2—fs4c | Passed | Two-way RM ANOVA followed by | Opsin x CS, $F_{(2,22)}$ = 1.926 | GFP vs. ChR2, BL/$CS^-$/$CS^+$, P > 0.9999, ns | GFP = 6 | NA |

**NA = Not applicable NS = Not significant fs = Figure supplement**

### Supplementary File 1 | Summary of statistical results associated with the Figures and figure supplements

| Figure number | Normality test | Statistic method | Main effect | Post hoc multiple comparisons test | Number of mice | Number of cells |
|---|---|---|---|---|---|---|
| | | Bonferroni post-hoc test | P = 0.1695, ns | | ChR2 = 7 | |
| 2—fs4d | Passed | Two-way RM ANOVA followed by Bonferroni post-hoc test | Opsin x CS, $F_{(2,22)}$ = 0.1444<br>P = 0.8663, ns | GFP vs. ChR2, BL/$CS^-$/$CS^+$, P > 0.9999, ns | GFP = 6<br>ChR2 = 7 | NA |
| 2—fs4e | Passed | Two-way RM ANOVA followed by Bonferroni post-hoc test | Opsin x CS, $F_{(2,22)}$ = 0.1095<br>P = 0.8967, ns | GFP vs. ChR2, BL/$CS^-$/$CS^+$, P > 0.9999, ns<br>BL vs. $CS^-$, GFP, P > 0.9999, ns<br>BL vs. $CS^-$, ChR2, P > 0.5328, ns<br>BL vs. $CS^+$, GFP/ChR2, P < 0.0001<br>$CS^-$ vs. $CS^+$, GFP/ChR2, P < 0.0001 | GFP = 6<br>ChR2 = 7 | NA |
| 2—fs5c | Failed | Friedman test followed by Dunn's multiple post-hoc test | P value < 0.0001<br>$Q$ = 19.85 | BL vs. $CS^-$, P > 0.9999, ns<br>BL vs. $CS^+$, P = 0.0012<br>$CS^-$ vs. $CS^+$, P = 0.0001 | 13 | NA |
| 2—fs5d | Passed | Two-way RM ANOVA followed by Bonferroni post-hoc test | Opsin x CS, $F_{(1,11)}$ = 12.39<br>P = 0.0048 | $CS^-$ vs. $CS^+$, GFP, P = 0.0001<br>$CS^-$ vs. $CS^+$, ChR2, P = 0.3342, ns<br>GFP vs. ChR2, $CS^-$, P > 0.9999, ns<br>GFP vs. ChR2, $CS^+$, P = 0.0300 | GFP = 6<br>ChR2 = 7 | NA |
| 2—fs5g | Passed | One-way RM ANOVA followed by Bonferroni post-hoc test | $F_{(2,18)}$ = 147.1<br>P < 0.0001 | BL vs. $CS^-$, P = 0.8655, ns<br>BL vs. $CS^+$, P < 0.0001<br>$CS^-$ vs. $CS^+$, P < 0.0001 | 10 | NA |
| 2—fs5h | Passed | Two-way RM ANOVA followed by Bonferroni post-hoc test | Opsin x CS, $F_{(1,8)}$ = 0.3984<br>P = 0.5455, ns | $CS^-$ vs. $CS^+$, GFP, P = 0.0001<br>$CS^-$ vs. $CS^+$, ArchT, P = 0.0003<br>GFP vs. ArchT, $CS^-$, P > 0.9999, ns<br>GFP vs. ArchT, $CS^+$, P = 0.3869, ns | GFP = 5<br>ArchT = 5 | NA |
| 2—fs6 | NA | No statistical test (anatomical verification of optic fiber placement) | NA | NA | 10 | NA |
| 4—fs1c | Passed | Paired t-test (two-tailed) | P = 0.6929, ns | $t_4$ = 0.4247 | 5 | 5 |
| 5—fs1b | A-β fiber, passed<br>A-δ fiber, passed<br>C fiber, passed | Paired t-test (two-tailed) | A-β fiber, P = 0.3750, ns<br>A-δ fiber, P = 0.7369, ns<br>C fiber, P = 0.4150, ns | A-β fiber, $t_6$ = 0.9564<br>A-δ fiber, $t_6$ = 0.3520<br>C fiber, $t_6$ = 0.8755 | 3 | A-β fiber, 7<br>A-δ fiber, 7<br>C fiber, 7 |

**NA = Not applicable NS = Not significant fs = Figure supplement**

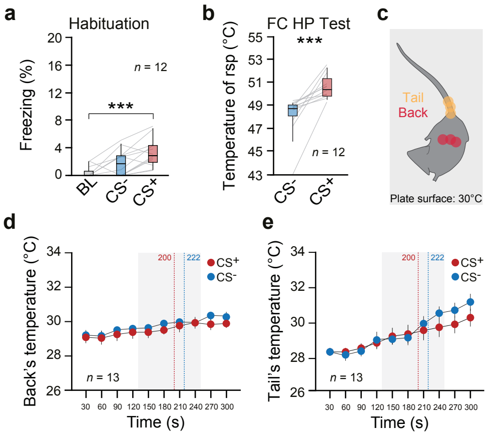
a
Habituation
Freezing (%)
n = 12
***
BL
CS-
CS+
b
FC HP Test
Temperature of rsp (°C)
***
n = 12
CS-
CS+
c
Tail
Back
Plate surface: 30°C
d
Back's temperature (°C)
200
222
CS+
CS-
n = 13
Time (s)
e
Tail's temperature (°C)
200
222
CS+
CS-
n = 13
Time (s)

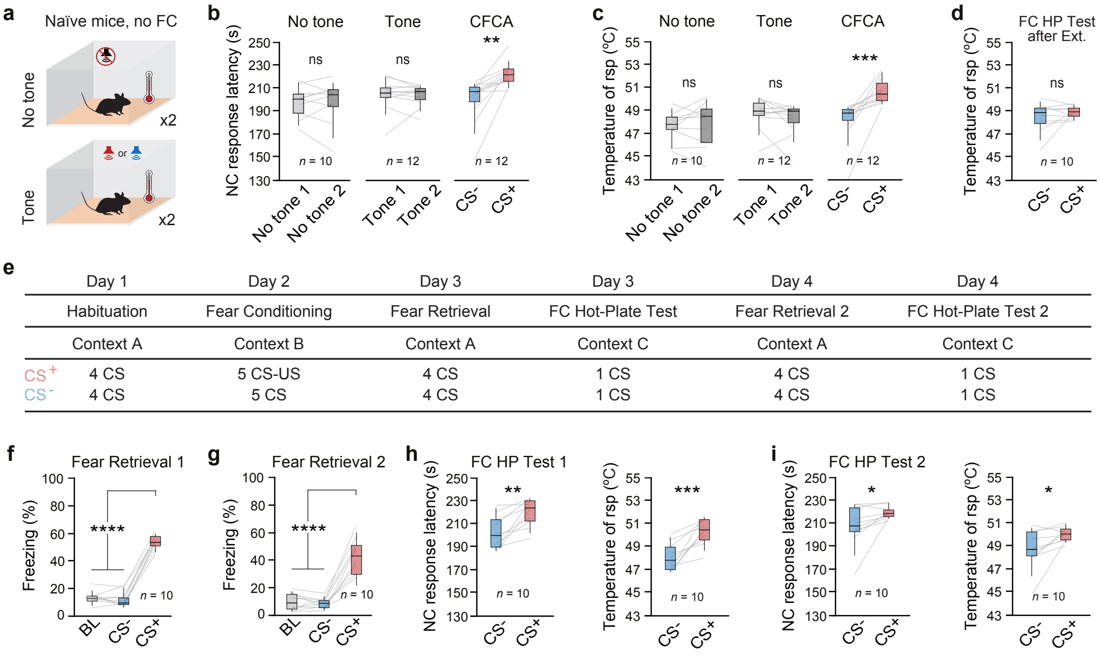
a
Naïve mice, no FC
No tone
x2
or
Tone
x2
b
No tone
Tone
CFCA
NC response latency (s)
ns
ns
**
n = 10
n = 12
n = 12
No tone 1
No tone 2
Tone 1
Tone 2
CS-
CS+
c
No tone
Tone
CFCA
Temperature of rsp (°C)
ns
ns
***
n = 10
n = 12
n = 12
d
FC HP Test after Ext.
Temperature of rsp (°C)
ns
n = 10
e
Day 1
Day 2
Day 3
Day 3
Day 4
Day 4
Habituation
Fear Conditioning
Fear Retrieval
FC Hot-Plate Test
Fear Retrieval 2
FC Hot-Plate Test 2
Context A
Context B
Context A
Context C
Context A
Context C
CS+
4 CS
5 CS-US
4 CS
1 CS
4 CS
1 CS
CS-
4 CS
5 CS
4 CS
1 CS
4 CS
1 CS
f
Fear Retrieval 1
Freezing (%)
****
n = 10
BL
CS-
CS+
g
Fear Retrieval 2
Freezing (%)
****
n = 10
h
FC HP Test 1
NC response latency (s)
**
n = 10
Temperature of rsp (°C)
***
n = 10
i
FC HP Test 2
NC response latency (s)
*
n = 10
Temperature of rsp (°C)
*
n = 10

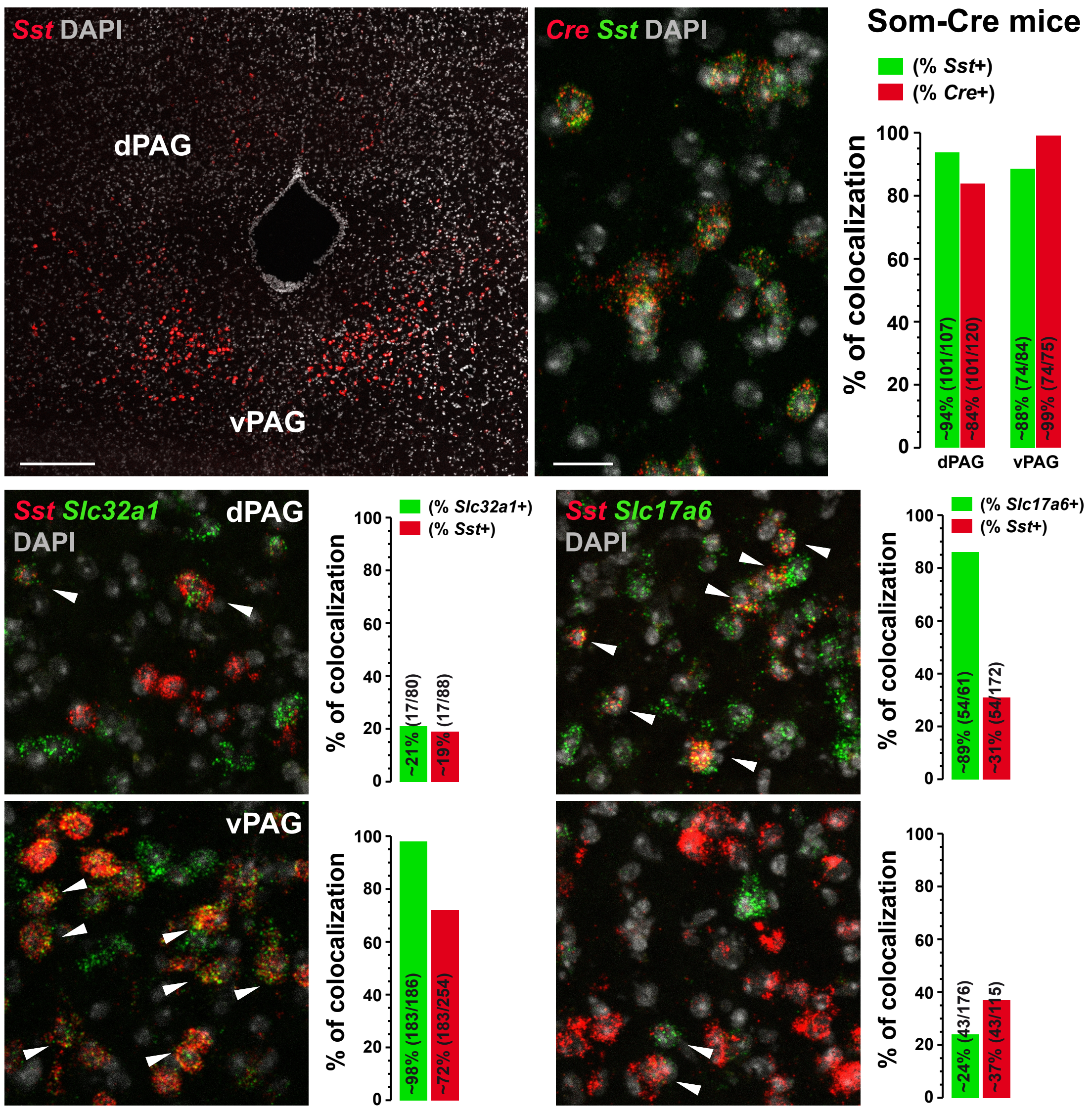

Sst DAPI
dPAG
vPAG
Cre Sst DAPI
Som-Cre mice
(% Sst+)
(% Cre+)
% of colocalization
~94% (101/107)
~84% (101/120)
~88% (74/84)
~99% (74/75)
dPAG
vPAG
Sst Slc32a1 DAPI
dPAG
(% Slc32a1+)
(% Sst+)
~21% (17/80)
~19% (17/88)
Sst Slc17a6 DAPI
(% Slc17a6+)
(% Sst+)
~89% (54/61)
~31% (54/172)
vPAG
~98% (183/186)
~72% (183/254)
~24% (43/176)
~37% (43/115)

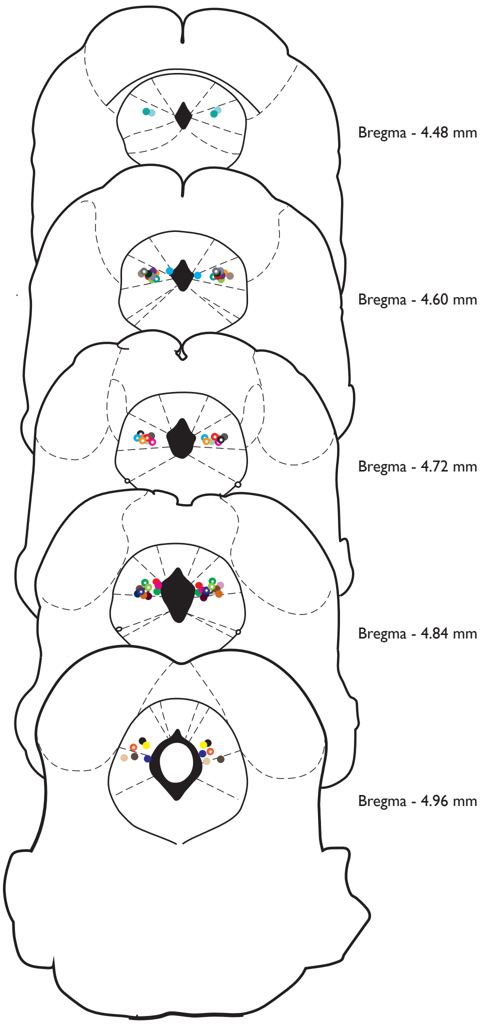
Bregma - 4.48 mm
Bregma - 4.60 mm
Bregma - 4.72 mm
Bregma - 4.84 mm
Bregma - 4.96 mm

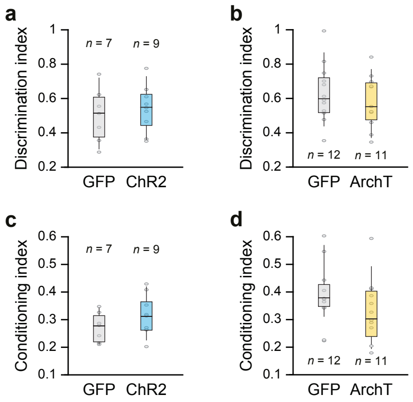
a
Discrimination index
1
0.8
0.6
0.4
0.2
n = 7
n = 9
GFP
ChR2
b
Discrimination index
n = 12
n = 11
GFP
ArchT
c
Conditioning index
0.6
0.5
0.4
0.3
0.2
0.1
n = 7
n = 9
GFP
ChR2
d
Conditioning index
n = 12
n = 11
GFP
ArchT

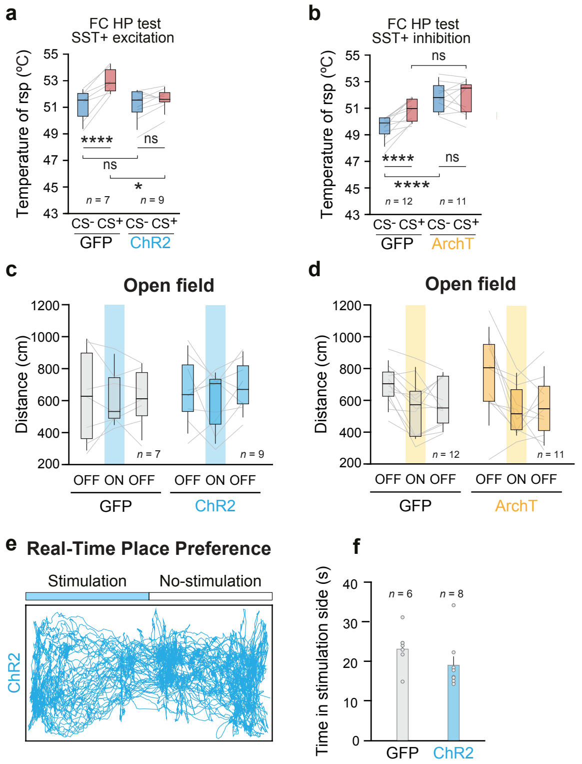

a
FC HP test
SST+ excitation
Temperature of rsp (ºC)
****
ns
ns
*
n = 7
n = 9
CS- CS+ CS- CS+
GFP ChR2
b
FC HP test
SST+ inhibition
Temperature of rsp (ºC)
ns
****
ns
****
n = 12
n = 11
CS- CS+ CS- CS+
GFP ArchT
c
Open field
Distance (cm)
n = 7
n = 9
OFF ON OFF OFF ON OFF
GFP ChR2
d
Open field
Distance (cm)
n = 12
n = 11
OFF ON OFF OFF ON OFF
GFP ArchT
e
Real-Time Place Preference
Stimulation
No-stimulation
ChR2
f
Time in stimulation side (s)
n = 6
n = 8
GFP ChR2

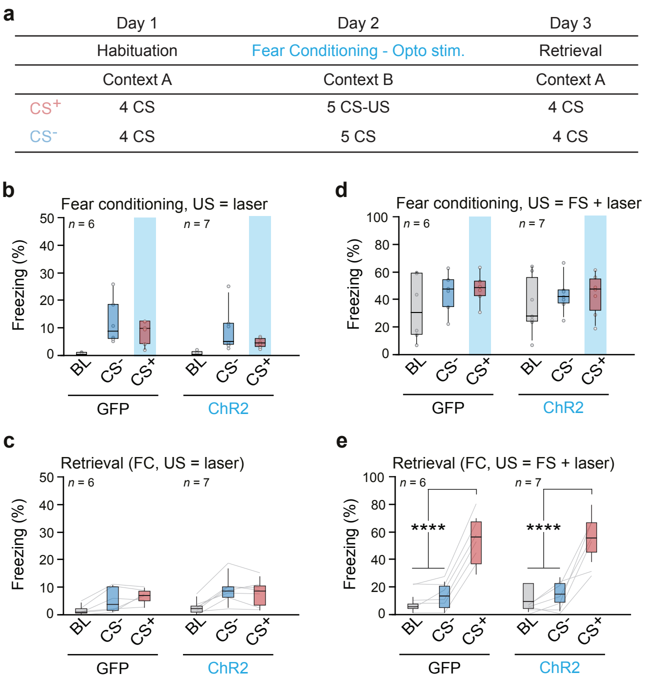

a
Day 1
Day 2
Day 3
Habituation
Fear Conditioning - Opto stim.
Retrieval
Context A
Context B
Context A
CS+
4 CS
5 CS-US
4 CS
CS-
4 CS
5 CS
4 CS
b
Fear conditioning, US = laser
Freezing (%)
n = 6
n = 7
BL
CS-
CS+
GFP
ChR2
c
Retrieval (FC, US = laser)
d
Fear conditioning, US = FS + laser
e
Retrieval (FC, US = FS + laser)
****
****

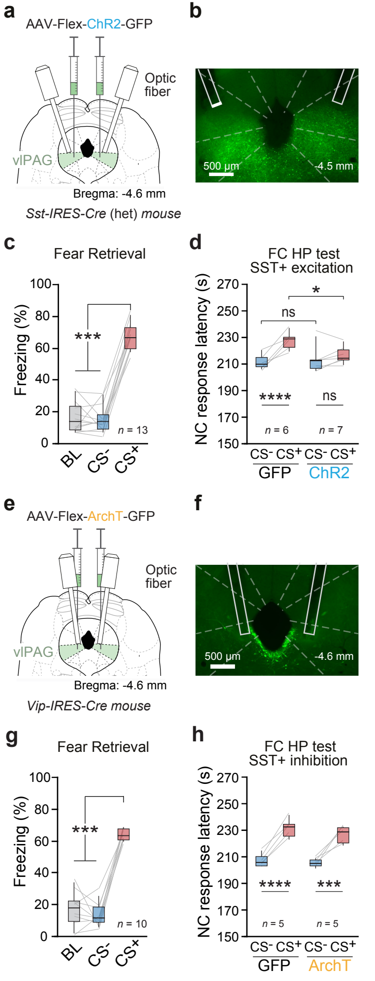

a
AAV-Flex-ChR2-GFP
Optic fiber
vlPAG
Bregma: -4.6 mm
Sst-IRES-Cre (het) mouse
b
500 µm
-4.5 mm
c
Fear Retrieval
Freezing (%)
***
n = 13
BL
CS-
CS+
d
FC HP test
SST+ excitation
NC response latency (s)
ns
*
****
ns
n = 6
n = 7
CS- CS+ CS- CS+
GFP
ChR2
e
AAV-Flex-ArchT-GFP
Optic fiber
vlPAG
Bregma: -4.6 mm
Vip-IRES-Cre mouse
f
500 µm
-4.6 mm
g
Fear Retrieval
Freezing (%)
***
n = 10
BL
CS-
CS+
h
FC HP test
SST+ inhibition
NC response latency (s)
****
***
n = 5
n = 5
CS- CS+ CS- CS+
GFP
ArchT

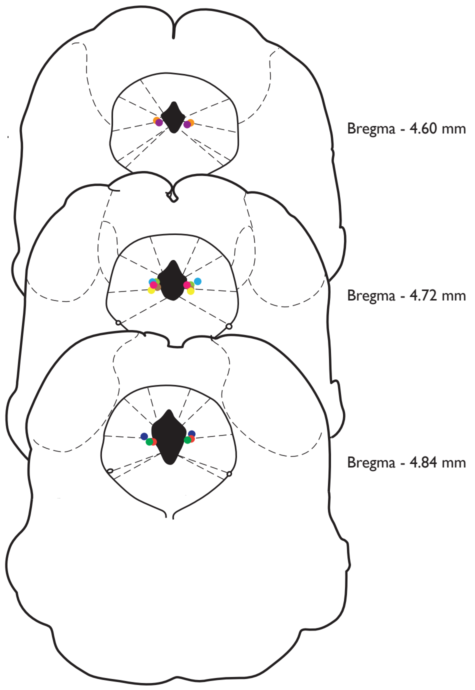
Bregma - 4.60 mm
Bregma - 4.72 mm
Bregma - 4.84 mm

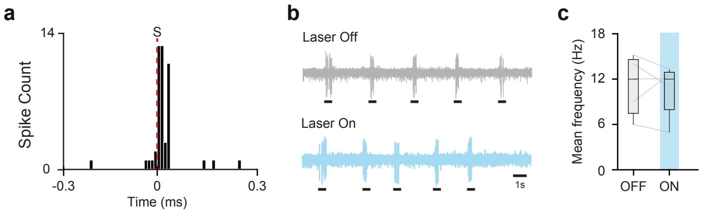

a
14
S
Spike Count
0
-0.3
0
0.3
Time (ms)
b
Laser Off
Laser On
1s
c
18
12
6
0
Mean frequency (Hz)
OFF
ON

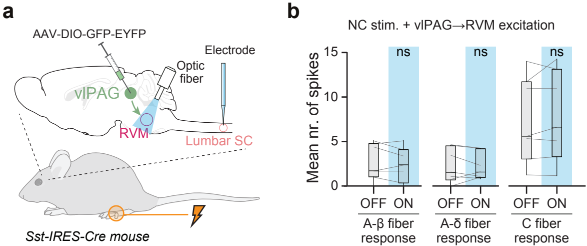
a
AAV-DIO-GFP-EYFP
vlPAG
Optic fiber
Electrode
RVM
Lumbar SC
Sst-IRES-Cre mouse
b
NC stim. + vlPAG→RVM excitation
Mean nr. of spikes
15
10
5
0
ns
ns
ns
OFF
ON
OFF
ON
OFF
ON
A-β fiber response
A-δ fiber response
C fiber response
